# c-Myc inhibits macrophage antimycobacterial response in *Mycobacterium tuberculosis* infection

**DOI:** 10.1101/2025.01.09.632095

**Authors:** Edoardo Sarti, Cédric Dollé, Rebekka Wolfensberger, Katharina Kusejko, Doris Russenberger, Simon Bredl, Roberto F. Speck, Melanie Greter, Jan H. Rueschoff, Lucas Boeck, Dat Mai, Alan H. Diercks, Peter Sander, Gregory S. Olson, Johannes Nemeth

## Abstract

*Mycobacterium tuberculosis* (MTB) is a major global cause of mortality worldwide, responsible for over a million deaths annually. Despite this burden, natural immunity prevents disease in more than 90% of exposed individuals. Previous studies have identified interferon-gamma (IFN-γ) as a key regulator of innate immune defense against MTB. Here, we investigate the impact of IFN-γ timing on macrophage-mediated control of MTB infection. We demonstrate that IFN-γ exposure before infection enhances macrophage antibacterial activity, whereas post-infection exposure does not. Further investigation into this phenotype revealed a strong association between c-Myc signaling and macrophage function in MTB control, as identified using unbiased in vitro systems approaches. Given the challenge of perturbing c-Myc in primary cells, we developed a lentiviral system for c-Myc inhibition and overexpression. Using a tetracycline-inducible Omomyc system - a small peptide inhibitor of c-Myc - we show that c-Myc inhibition promotes a pro-inflammatory macrophage phenotype with enhanced antimycobacterial activity. Mechanistically, c-Myc inhibition induces metabolic reprogramming via increased mTORC1 activity, leading to upregulated inducible nitric oxide synthase and improved bacterial control. In vivo analyses, including murine models and human clinical histopathology, reveal a strong correlation between c-Myc expression and MTB persistence, as well as active tuberculosis (TB), suggesting a role for c-Myc in immune evasion. These findings reveal c-Myc as a potential mediator of immune privilege in MTB infection and highlight its role as a promising target for novel TB therapies aimed at enhancing macrophage function.

## Introduction

*Mycobacterium tuberculosis* (MTB) remains the leading cause of death from a single infectious agent worldwide, responsible for approximately 1.25 million deaths annually, according to the World Health Organization (WHO)^1^. The emergence and spread of antibiotic-resistant strains of MTB demand alternative tools beyond antibiotics to achieve the objectives of global tuberculosis (TB) control efforts.

Although MTB infection is either cleared or effectively controlled by the host immune system in most cases, no host-directed therapies exist for the treatment of active TB^2^. Observations from animal models, human studies, natural infections with HIV-1, and new anti-inflammatory therapies (e.g. anti-TNF therapy) have identified pro-inflammatory macrophages and T cells to be critical components of an effective host immune response^3^. The ensuing immune response may result in profound heterogeneity in the outcomes of MTB infection^4^, ranging from deadly disease to complete clearance without any sequelae^2,5,6^. This variability is observed not only across populations but also within individual infected organisms across different species, including mice, non-human primates, and humans^6–8^.

To investigate competent host immunity, we have previously focused on asymptomatic MTB infection, the most common, yet underexplored, outcome of MTB infection. Asymptomatic infection appears to confer protection against reinfection - a phenomenon observed in mouse models, non-human primates, and humans^9–11^. In our studies, mice initially received an intradermal inoculation, leading to an asymptomatic primary infection localized within the draining lymph node (contained MTB infection, CMTB). Compared to uninfected controls, mice with CMTB demonstrated a significantly faster and stronger immune response to a secondary, pulmonary challenge. This response included heightened activation of alveolar macrophages (AMs) and rapid recruitment of inflammatory monocytes and T cells, driven by low-grade systemic interferon (IFN)-γ. Consequently, these mice exhibited a substantially lower bacterial burden in the lungs than those without prior infection^9^.

IFN-γ is a key cytokine in both innate and adaptive immunity, primarily produced by NK cells, CD4^+^ T cells, and CD8^+^ T cells^12^. Under stress or infection, IFN-γ enhances macrophage activation by increasing proinflammatory cytokines and suppressing anti-inflammatory responses, which supports both innate and adaptive immunity^13,14^. It also boosts macrophage phagocytic and cytolytic abilities, facilitating pathogen elimination - a process termed classical macrophage activation. Additionally, IFN-γ triggers signaling cascades that upregulate genes for reactive oxygen and nitrogen species, while also influencing metabolic pathways to meet the energy demands of activated macrophages in response to external stressors, including mycobacterial infections^13^.

Interestingly, in cases of active TB in humans, there are often high levels of IFN-γ present at the infection site^15,16^. This counterintuitive observation suggests that, despite the abundance of this potent pro-inflammatory cytokine, certain tissues may not be responding adequately to the IFN-γ stimulus^4^. This impaired response could indicate a form of immune resistance or dysfunction^17^, where macrophages and other immune cells fail to fully activate or sustain their antimicrobial functions even in the presence of IFN-γ^2,3,18^.

Based on this conundrum, the starting point for the following work was the hypothesis that the timing of IFN-γ exposure on macrophages relative to MTB infection is crucial for their effective response to MTB infection. Thus, we hypothesized that the sequence of cytokine exposure and MTB infection could significantly influence macrophage activation states and, consequently, their ability to control bacterial growth. Therefore, we decided to investigate how the timing of IFN-γ exposure - whether before or after in vitro infection - affects macrophage activation, gene expression, and antibacterial activity.

## Results

### The sequence of infection and activation determines the outcome of *Mycobacterium tuberculosis* infection in macrophages

We hypothesized that the timing of IFN-γ exposure is a critical factor in determining infection outcomes. To test this, we conducted an in vitro experiment using C57BL/6 murine bone marrow-derived macrophages (BMDMs) divided into four groups. To assess bacterial growth, we used a colony-forming unit (CFU) assay, in which macrophages were lysed, and cell lysates were plated on Middlebrook 7H10 agar to quantify viable MTB by counting bacterial colonies.

The experimental groups included: (1) macrophages continuously exposed to IFN-γ throughout the 6-day experiment of infection with the virulent MTB strain H37Rv, including the pretreatment period, (2) macrophages pre-treated with IFN-γ starting 24 hours prior to infection, but thereafter removed, (3) macrophages treated with IFN-γ immediately post-infection (approximately 1 hour after phagocytosis) and maintained for 5 days, and (4) a control group with no IFN-γ treatment (Figure 1A).

**Figure 1.**
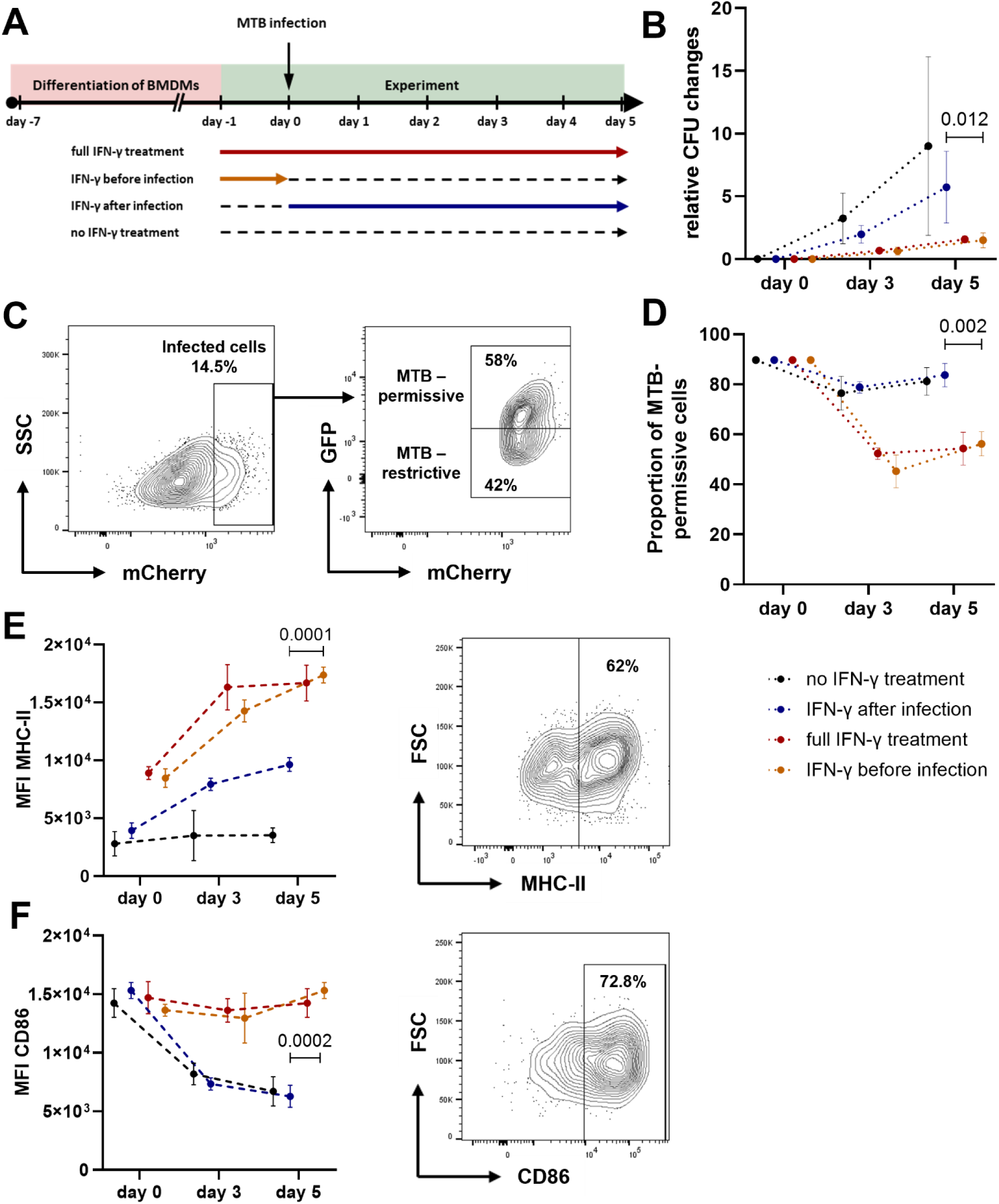
Antimycobacterial properties of IFN-γ depend on the sequence of infection and IFN-γ exposure. **A)** Experimental workflow. BMDMs were isolated from C57BL/6 mice and differentiated for 6 days. Cells were either pre-activated with IFN-γ (25 ng/mL) 24 h before infection or treated immediately after infection. Infection was performed with MTB H37Rv (MOI 5). **B)** Kinetics of intracellular bacterial burden. CFU/mL of intracellular MTB H37Rv were measured immediately after phagocytosis (day 0) and at days 3 and 5 post-infection. Data are mean ± SD of five independent experiments, normalized to day 0. **C)** A representative flow cytometry plot showing viable F4/80⁺CD11b⁺ BMDMs infected with MTB H37Rv live-dead strain is shown. Infected (mCherry⁺) cells are subdivided into “permissive” or “restrictive” populations based on GFP induction (1 µg/mL doxycycline) after 24 h. **D)** Percentage of “permissive” macrophages (GFP⁺) at days 0, 3 and 5 post-infection. **E–F)** Activation marker expression. Median fluorescence intensity (MFI) of MHC-II (E) and CD86 (F) in infected BMDMs at days 0, 3 and 5, with representative plots (full-treatment condition shown). Unless otherwise stated, data are mean ± SD (n = 3). Statistical analysis: *P* values were determined by unpaired two-tailed Student’s t-test (before vs. after infection) and corrected for multiple testing using the two-stage linear step-up method of Benjamini–Krieger–Yekutieli.

Results from CFU assays revealed that only macrophages exposed to IFN-γ prior to infection, with or without continued exposure, showed improved control of MTB growth. In contrast, macrophages treated with IFN-γ post-infection did not exhibit enhanced antimycobacterial activity, as indicated by higher CFU counts (Figure 1B + Supplementary Figure 1).

To assess both functionality and survival of phagocyted MTB, as well as the transcriptional state at the single-cell level, we utilized the MTB live-dead strain^19^. This strain constitutively expresses mCherry, enabling identification of cells containing phagocytosed bacteria, and expresses GFP under the control of a tetracycline-inducible promoter. Bacteria capable of activating GFP expression in response to doxycycline (DOX), a synthetic version derived from tetracycline, are defined as transcriptionally active (mCherry⁺/GFP⁺), while those that do not are considered inactive (mCherry⁺/GFP⁻) (Figure 1C).

While the live-dead strain has primarily been used in fluorescence microscopy^19–21^, we developed a new application of this tool in flow cytometry (Supplementary Figure 2). To validate this approach, we compared the live-dead strain results with CFU assays using isoniazid-treated MTB H37Rv, finding a comparable reduction in transcriptional activity and CFU counts (Supplementary Figure 3).

Using flow cytometry and the live-dead strain, we confirmed that macrophages exposed to IFN-γ prior to infection suppress transcriptional activity in approximately 50% of intracellular MTB. In contrast, macrophages treated with IFN-γ post-phagocytosis were unable to inhibit MTB transcriptional activity (Figure 1D).

Simultaneously, this assay allowed us to evaluate macrophage activation, with enhanced antimycobacterial activity correlating with upregulation of MHC-II, CD86 and CD80 surface markers (Figures 1E-1F + Supplementary Figure 4). These results suggest that the increased bacterial control observed in CFU assays was due to the heightened activation of viable BMDMs, rather than an artifact from macrophage death or extracellular bacteria.

### c-Myc associated transcriptional programs mirror the in vitro phenotype in mature macrophages

To assess the transcriptome underlying the different infection timepoints and activation states, we conducted bulk RNA sequencing (RNA-seq) at 6 hours and 24 hours post-infection in the previously described groups of BMDMs. Principal Component Analysis (PCA) revealed that as early as 6 hours post-infection, macrophages activated prior to infection began clustering distinctly from those activated after infection (Figure 2A). Specifically, macrophages activated before infection clustered more closely with fully treated cells (MTB-controlling cells), while those activated post-infection clustered nearer to untreated cells (MTB-permissive cells). This distinction became more pronounced at 24 hours, with macrophages activated prior to infection clustering with fully treated cells, whereas those activated post-infection formed a separate cluster, distinct from both fully activated and untreated cells.

**Figure 2:**
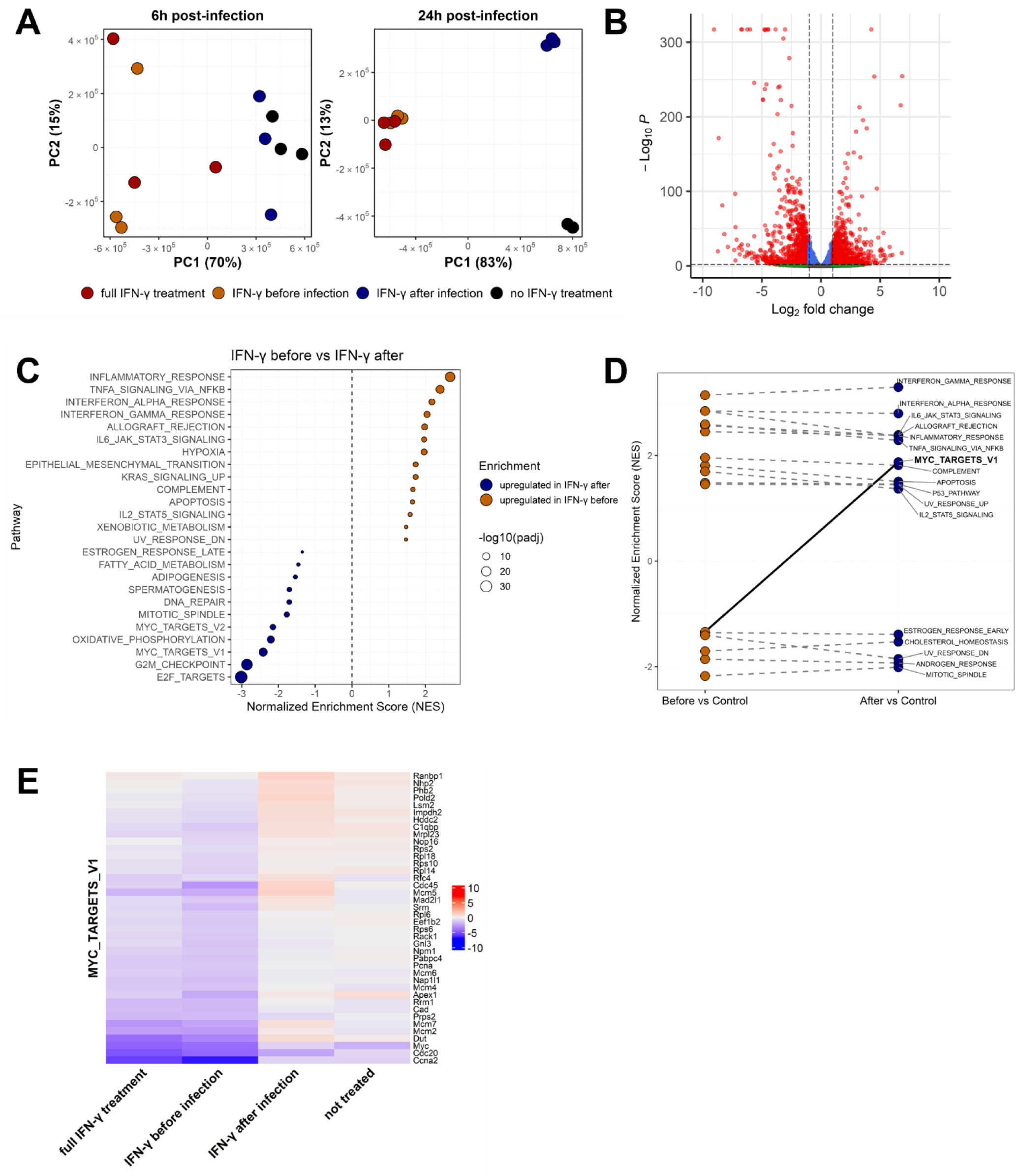
Changes in c-Myc associated transcriptional programs mirror MTB permissive versus controlling macrophages in vitro. **A)** Principal component analysis (PCA) of differently activated/infected BMDMs, 6 and 24 h after infection with MTB H37Rv (MOI 5). Each point represents an individual sample, with colors indicating the treatment. PC1 and PC2 are plotted with the percent variance explained indicated. **B)** Volcano plot of differential expression in BMDMs 24 h post-infection, comparing IFN-γ after vs. IFN-γ before conditions. Genes are plotted by log_2_ fold-change (x-axis) against –log_10_(adjusted *P*) (y-axis). Red dots mark genes meeting |log_2_FC| > 2 and adjusted *P* < 0.01; dashed lines denote these thresholds. **C)** Gene set enrichment analysis (GSEA) dot plot of Hallmark pathways from BMDMs 24 h post-infection, comparing IFN-γ administered before infection versus IFN-γ added after infection. Dots represent normalized enrichment scores (adjusted *P* < 0.05), with positive values indicating enrichment in the before condition. NES values were computed by fgsea on MSigDB Hallmarks (v2024.1.Mm). **D)** Connected-dot plot of GSEA comparing IFN-γ–activated BMDMs before and after relative to untreated, uninfected controls. Each point represents one gene set’s NES: orange circles mark sets significantly enriched in the before vs. control comparison (adjusted *P* < 0.05), blue circles mark sets significantly enriched in the after vs. control comparison (adjusted *P* < 0.05). The y-axis shows NES values (positive = up-regulation; negative = down-regulation). Gray dashed lines connect the same gene set’s NES across the two contrasts; the solid black line highlights those gene sets that reverse their direction of regulation between pre- and post-infection. **E)** Heatmap of leading-edge genes (top 40 genes) of the MYC_TARGETS_V1 set across all four conditions 24 h after infection. The differential expression is calculated as mean log_2_ fold-change per condition compared to untreated, uninfected BMDMs (n = 3). All data shown were derived from bulk RNA sequencing.

These findings indicate that as early as 6 hours post-infection, macrophages begin to display trajectories predictive of the infection outcome. The observation that cells receiving IFN-γ post-infection diverge from unstimulated cells but do not cluster with cells that received IFN-γ prior to infection suggests that while these cells can respond to IFN-γ, their trajectory differs significantly from those pre-activated with IFN-γ.

At the single gene level, macrophages treated before infection and macrophages treated after infection have a clearly different expression signature, with 311 significantly upregulated genes and 396 significantly downregulated genes (Figure 2B).

We performed Gene Set Enrichment Analysis (GSEA) for functional categorization. Comparisons of gene expression profiles between macrophages activated prior to infection versus macrophages activated after infection revealed a relative downregulation of multiple pro-inflammatory pathways in cells that received IFN-γ after infection (Figure 2C). Notably, however, the most significant changes, in terms of fold changes, were not among the classical pro-inflammatory genes but were instead related to cell cycle genes, including E2F targets, G2M targets, and MYC-associated genes, which are upregulated in macrophages treated after infection (Supplementary Figure 5). Additional comparisons of pre- and post-treated conditions with untreated, uninfected BMDMs confirmed an activation of IFN-γ–responsive pathways in both treated conditions, indicating that IFN-γ elicited a strong transcriptional response regardless of timing (Supplementary Figure 6). Focusing on all significantly up- and downregulated pathways, it is striking that only MYC-associated genes switch regulation between pre- and post-treatment conditions (Figure 2D).

Among all transcriptional regulators and pathways analyzed, c-Myc was the only transcription factor whose associated gene expression profile directly mirrored the in vitro antimycobacterial capacity of macrophages. Specifically, MYC-associated genes were the only gene set that fully recapitulated the phenotype distinguishing MTB-permissive from MTB-controlling cells, and thus represent a potentially targetable axis. Macrophages that effectively restricted MTB replication exhibited downregulation of leading-edge genes in the c-Myc-associated pathway (Figure 2E), suggesting that suppression of MYC activity is linked to enhanced antimycobacterial function. By contrast, cells stimulated with IFN-γ after infection showed transcriptional evidence of IFN-γ signaling but did not demonstrate improved mycobacterial control (Figure 1), indicating that this response alone was insufficient to drive effective restriction of MTB.

These findings suggest that at the transcriptional level, the ability to downregulate c-Myc-associated transcriptional programs is strongly associated with a macrophage’s capacity to mount an effective immune response that controls in vitro MTB infection.

### c-Myc inhibition leads to a gain of antimycobacterial function via a distinct transcriptional program

Based on the strong associations reported above, we decided to experimentally perturb c-Myc in macrophages to better understand its role in mycobacterial infections.

To achieve this, we developed a lentiviral system featuring constitutive expression of a truncated nerve growth factor receptor (NGFR) and tetracycline (Tet)-inducible GFP (Supplementary Figure 7). This system was used to transduce bone marrow cells from C57BL/6 mice. After successful transduction, cells were magnetically sorted and differentiated into BMDMs. When exposed to doxycycline these cells expressed GFP, confirming the functionality of our inducible system (Supplementary Figure 8).

Next, we replaced GFP with murine c-Myc or Omomyc, a small peptide inhibitor of c-Myc^22,23^. RNA-seq analysis then confirmed that, as expected, all transduced cells expressed the truncated NGFR transduction marker (Figure 3A).

**Figure 3:**
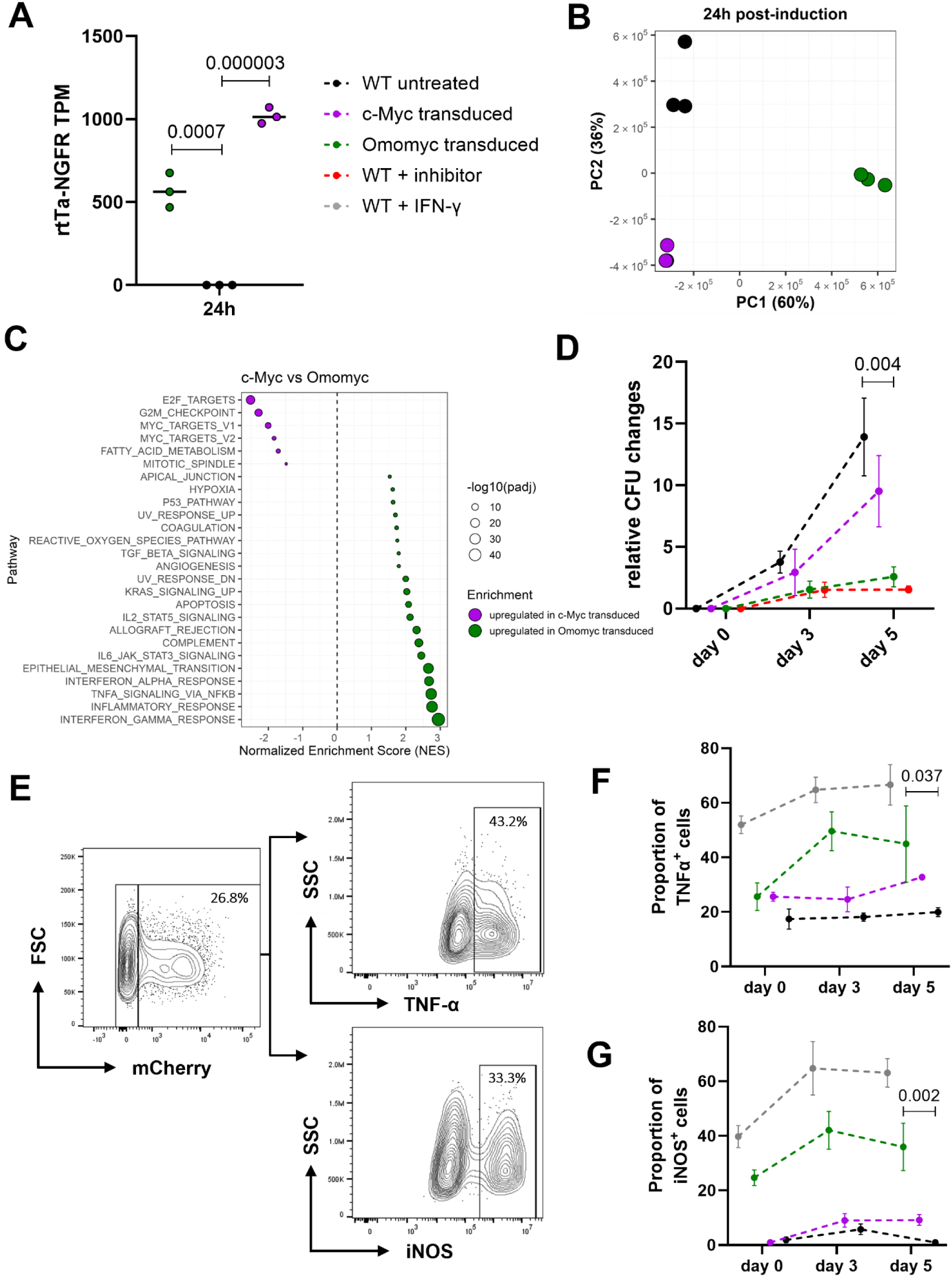
c-Myc inhibition leads to a gain in anti-mycobacterial function and induces a pro-inflammatory phenotype in macrophages in vitro. **A)** RNA expression of rtTa-NGFR (transduction marker) was measured by bulk RNA sequencing as transcripts per million (TPM) in transduced BMDMs after 24 h (n = 3). **B)** Principal component analysis (PCA) of BMDMs expressing either c-Myc or Omomyc versus WT 24 h after Tet-on induction with doxycycline (100ng/mL). Each point represents an individual sample, with colors indicating the experimental group. PC1 and PC2 are plotted with the percent variance explained on the axes. **C)** Gene set enrichment analysis (GSEA) dot plot of Hallmark pathways from BMDMs expressing Omomyc compared to BMDMs expressing c-Myc 24 h after Tet-on induction with doxycycline. Dots represent normalized enrichment scores (adjusted *P* < 0.01), with positive values indicating enrichment in the Omomyc group. NES values were computed by fgsea on MSigDB Hallmarks (v2024.1.Mm). **D)** Kinetics of intracellular bacterial burden in WT BMDMs, BMDMs treated with the chemical c-Myc inhibitor 10058-F4, and BMDMs expressing c-Myc or Omomyc. CFU/mL of intracellular MTB H37Rv were measured immediately after phagocytosis (day 0) and at days 3 and 5 post-infection. Data are mean ± SD of three independent experiments, normalized to day 0. **E)** Representative flow cytometry plots of fully mature BMDMs infected with mCherry-expressing MTB, gated on mCherry⁺ cells, showing intracellular TNF-α and iNOS expression (Omomyc-transduced condition is shown). **F–G)** Quantification of the proportion of mCherry⁺ BMDMs expressing (F) TNF-α and (G) iNOS at days 0, 3 and 5 post-infection. WT BMDMs, WT pre-treated with IFN-γ, and BMDMs expressing c-Myc or Omomyc were infected and analyzed by flow cytometry. Data are mean ± SD of three independent experiments. Statistical analysis: *P* values were determined by unpaired two-tailed Student’s t-test (Omomyc transduced vs. WT untreated) and corrected for multiple testing using the two-stage linear step-up method of Benjamini–Krieger–Yekutieli.

To ensure that c-Myc perturbation did not disrupt macrophage function, we assessed phagocytic activity and phagosome acidification in transduced and untransduced BMDMs. We used pHrodo Green *E. coli* bioparticles and mCherry-expressing MTB H37Rv. pHrodo Green is a pH-sensitive fluorescent dye that becomes increasingly fluorescent in acidic environments, allowing selective detection and quantification of active phagocytosis and phagosome acidification^24^. No significant differences were observed between Omomyc-transduced and wildtype macrophages in either the extent of phagocytosis or phagosome acidification (Supplementary Figure 9).

With the system validated, we exposed the transduced BMDMs (Supplementary Figure 10) to doxycycline for 24 hours and performed RNA-seq. The transcriptomic analysis revealed that both c-Myc overexpression and Omomyc-mediated inhibition had profound effects on the cells’ transcriptome (Figure 3B). While the significant changes induced by c-Myc overexpression were anticipated - given c-Myc’s established role in reprogramming somatic cells into induced pluripotent stem cells^25^ - it was particularly striking that Omomyc inhibition also caused extensive transcriptomic alterations.

Subsequent functional enrichment analysis using GSEA indicated that Omomyc-expressing cells exhibited a distinct transcriptional response (Figure 3C). Notably, these cells showed a pronounced upregulation of inflammatory responses such as IFN-γ, tumor necrosis factor (TNF)-α or IFN-α response, and a general increase of metabolic associated pathways, such as an increase of the reactive oxygen species pathway. These findings suggests that c-Myc plays a critical role in the regulation of gene expression in mature macrophages in the short term (24 hours), and its up- or downregulation is sufficient to induce changes both in key inflammatory transcriptional responses and the metabolism in macrophages.

To directly assess how c-Myc inhibition impacts MTB growth control, we infected Omomyc- and c-Myc-expressing macrophages with MTB H37Rv. Based on our previous data, we hypothesized that c-Myc inhibition would enhance the macrophages’ ability to control MTB growth. Indeed, Omomyc-expressing macrophages demonstrated that c-Myc inhibition led to a significant gain of antimycobacterial activity, comparable to the effect observed with IFN-γ pretreatment (Figure 3D). As an additional control, we tested a commercially available chemical c-Myc inhibitor, 10058-F4, which also led to an increase in antimycobacterial activity, confirming our findings. At the same time, the construct expressing c-Myc had no significant effect on MTB growth. In order to exclude possible antibiotic side effects of doxycycline on bacterial growth, we also compared all tested conditions with and without the addition of doxycycline, thus also eliminating any overexpression of Omomyc and c-Myc and detected no differences in MTB H37Rv growth depending on the presence of doxycycline, as was also shown in other studies^26^ (Supplementary Figure 11). Finally, the addition of IFN-γ showed that all conditions responded with similar dynamics in bacterial growth when pre-treated, further demonstrating that overexpression or inhibition of the key transcription factor c-Myc is tolerated in the presented system (Supplementary Figure 12).

To understand how these transcriptional changes impacted the macrophages at a cellular level over time, we infected BMDMs with mCherry-expressing MTB H37Rv (Figure 3E) and monitored key functional markers by flow cytometry. As predicted by the transcriptional profile, the Omomyc-expressing macrophages displayed an upregulation of TNF-α on the protein level (Figure 3F). Also, the cells mounted a robust inducible nitric oxide synthase (iNOS) response (Figure 3G). These findings indicated that the macrophages had gained enhanced antimycobacterial function, consistent with the prediction based on transcriptional changes. Importantly, the Omomyc mediated inhibition of c-Myc did not significantly affect macrophage survival over time compared to control cells as measured with cell viability staining and cell counting, suggesting that the observed changes were specific to the modulation of c-Myc activity rather than a general cytotoxic effect (Supplementary Figure 13).

In conclusion, our data demonstrate that c-Myc is a crucial regulator of macrophage function after MTB infection in vitro. Inhibition of c-Myc via Omomyc induces a unique transcriptional profile that enhances the antimycobacterial activity of macrophages.

### mTORC1 mediated metabolic programming partially explains enhanced antimycobacterial function in c-Myc-inhibition in macrophages

We observed that inhibition of c-Myc via the Omomyc peptide led to significant changes in macrophage functionality, notably an enhancement in antimycobacterial activity. This finding prompted us to investigate the underlying mechanisms, particularly focusing on metabolism. iNOS, which is essential for the production of nitric oxide (NO), a key antimicrobial effector^27^, requires a substantial energy investment, suggesting a need for metabolic reprogramming to support this heightened immune response. Considering the extensive evidence on c-Myc’s central role in cellular metabolism, we postulated that c-Myc inhibition might facilitate a metabolic shift that supports enhanced macrophage function. Our transcription analysis supported this hypothesis and showed a significant increase in several metabolic pathways in c-Myc-inhibited macrophages, including mTORC1 signaling (Figure 4A-B). Therefore, we proposed that the gain of function observed through c-Myc inhibition is driven by metabolic changes mediated by mTORC1.

**Figure 4:**
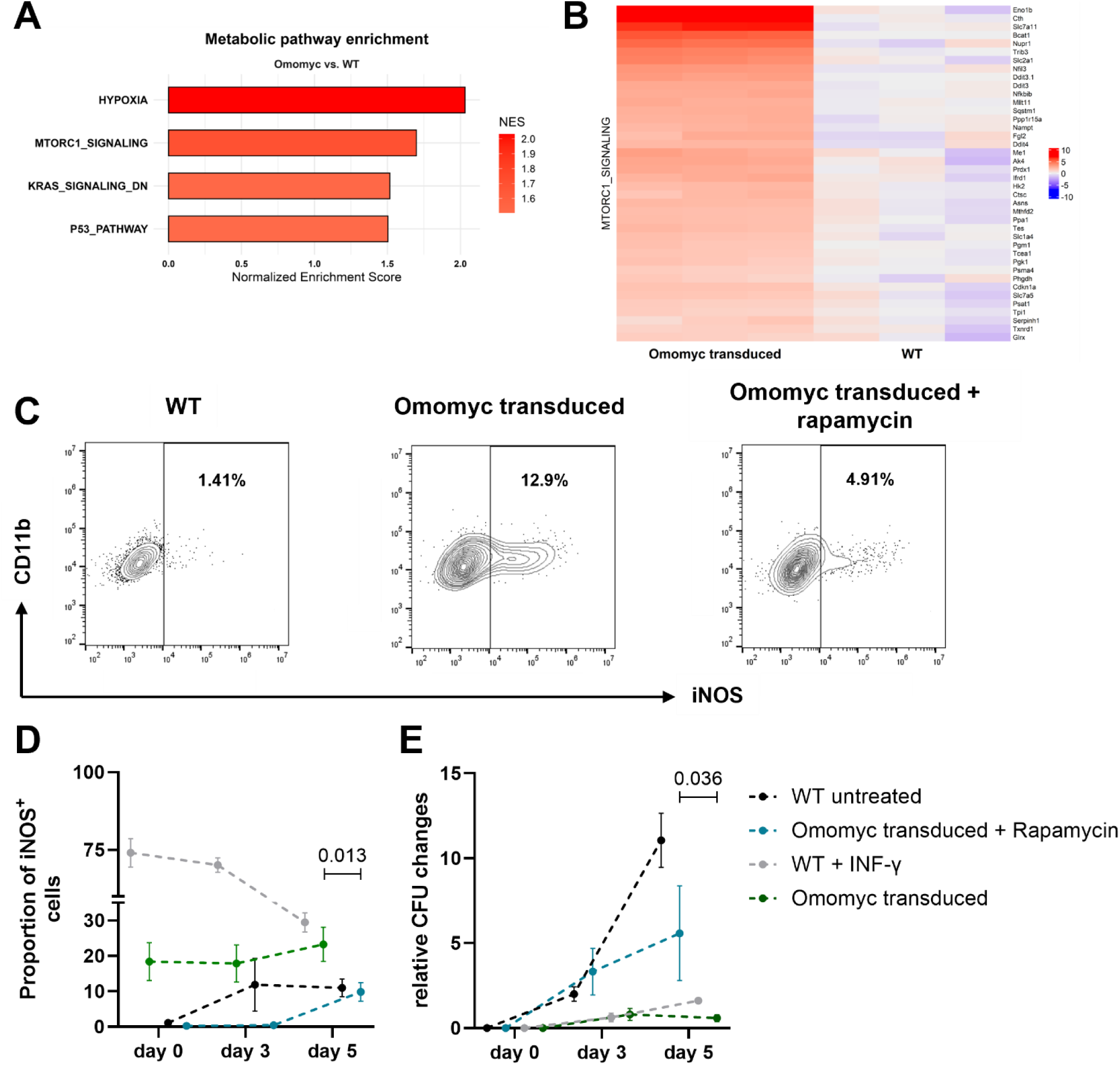
c-Myc inhibition associated gain of antimycobacterial function is partly mediated by a shift towards mTORC1 signaling. **A)** Bar plot of Hallmark metabolic pathways significantly enriched by gene set enrichment analysis (GSEA) (adjusted *P* < 0.01). Bars show normalized enrichment scores (NES), positive NES (red bars) denote enrichment in the Omomyc group. **B)** Heatmap of leading-edge genes from the MTORC1_SIGNALING Hallmark pathway. Rows are genes contributing most to the enrichment score, columns are individual Omomyc-expressing or WT BMDM samples. Values are log_2_ fold-change relative to the average of WT samples. **C)** Representative flow cytometry plots showing the proportion of iNOS⁺ cells in mCherry⁺ BMDMs on day 3 after infection with mCherry-expressing MTB (MOI 5). WT cells, Omomyc-expressing cells and Omomyc-expressing cells treated with 10 µM rapamycin are displayed. All samples received 0.1% DMSO. **D)** Proportion of infected (mCherry⁺) BMDMs expressing iNOS at days 0, 3 and 5 post-infection in WT, Omomyc-expressing, Omomyc-expressing + rapamycin, and WT + IFN-γ conditions. Data are mean ± SD (n = 3). **E)** Kinetics of intracellular bacterial burden. CFU/mL of MTB H37Rv were measured in WT BMDMs, Omomyc-expressing BMDMs, Omomyc-expressing + rapamycin, and WT + IFN-γ BMDMs immediately after phagocytosis (day 0) and at days 3 and 5 post-infection. Data represent mean relative changes (range) ± SD from three independent experiments, normalized to day 0. Statistical analysis: *P* values were determined by unpaired two-tailed Student’s t-test (Omomyc transduced vs. Omomyc transduced + rapamycin) and corrected for multiple testing using the two-stage linear step-up method of Benjamini–Krieger–Yekutieli.

To investigate this, we conducted in vitro infections using mCherry-expressing MTB H37Rv and measured iNOS production with or without mTORC1 inhibition via rapamycin. Prior to these experiments, we determined a concentration of rapamycin that effectively blocked the mTORC1 pathway without impairing macrophage viability (Supplementary Figure 14). In line with our hypothesis, rapamycin treatment significantly reduced Omomyc-induced iNOS production (Figure 4C-D), supporting the idea that mTORC1 plays a key role in the increased iNOS expression seen in c-Myc-inhibited BMDMs. We then infected BMDMs with MTB H37Rv. As expected, inhibiting mTORC1 reversed in part the enhanced antimycobacterial function caused by c-Myc inhibition (Figure 4E). This suggests that the improved antimycobacterial activity in c-Myc-inhibited macrophages is partially due to mTORC1 driven changes in metabolism.

Overall, these findings indicate that c-Myc inhibition leads to metabolic reprogramming including mTORC1, resulting in enhanced macrophage antimicrobial activity.

### c-Myc expression in myeloid cells is associated with MTB persistence in the contained MTB infection model

Given the central role c-Myc played in in vitro infections of BMDMs, we aimed to determine whether similar signals could be observed in vivo. To this end, we employed the CMTB model to investigate an in vivo system of asymptomatic MTB infection. As published previously, mice were intradermally infected with MTB H37Rv and allowed to develop a chronic infection over a six-week period^9^. After this time, lymph nodes were harvested, and B and T cells were removed via positive selection using magnetic beads. The remaining cells were subjected to single-cell RNA sequencing (scRNA-seq) using the 10X Genomics platform, with subsequent deconvolution of cell populations performed using the Seurat package (Figure 5A + Supplementary Figure 15) as published previously^28^. We observed a substantial increase of monocytes in infected lymph nodes (Supplementary Figure 16). Functional enrichment analysis focused on monocytes and macrophages, utilizing the AddModuleScore function to calculate the average expression levels of GSEA hallmark c-Myc-related gene programs at the single-cell level (Figure 5B).

**Figure 5:**
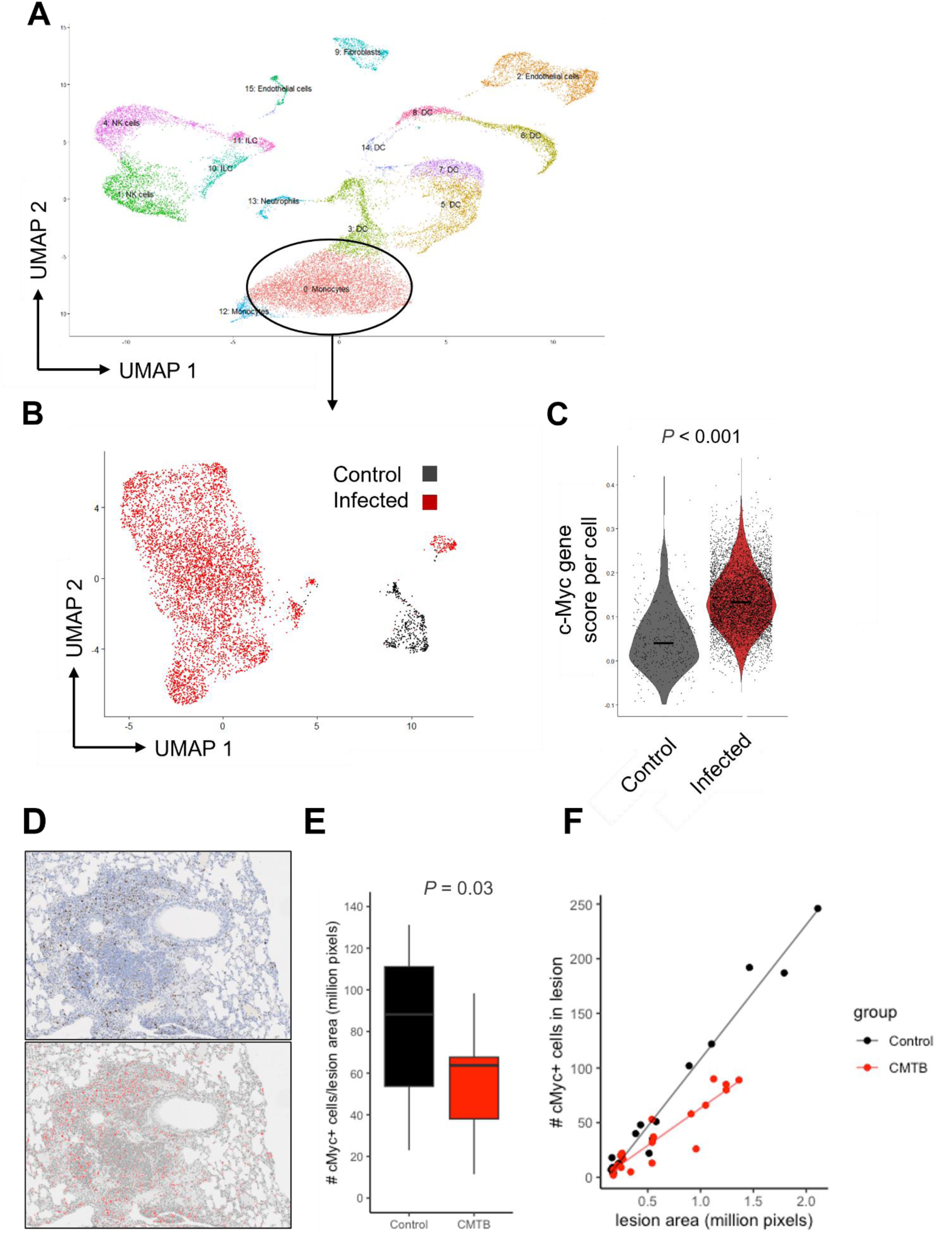
c-Myc expression is associated with MTB infection in the contained MTB infection (CMTB) mouse model of tuberculosis and in the pulmonary infection model. **A)** Lymph nodes from control and CMTB mice were depleted of CD3⁺/CD20⁺ cells and subjected to scRNA-seq. Major clusters were annotated by ImmGen matching, with the monocyte/macrophage cluster circled. **B)** UMAP of the circled monocyte/macrophage cluster showing clear segregation of infected versus non-infected cells. **C)** Single-cell c-Myc gene scores in infected versus control cells, calculated using AddModuleScore (aggregated expression of control gene set subtracted). **D)** Representative immunohistochemistry pathology slides (previously published^9^) from mice with and without CMTB and stained with an anti c-Myc antibody. Lesion size and number of c-Myc-expressing cells were analyzed using a semiautomated pipeline. **F)** Boxplot showing the density of c-Myc-expressing cells measured as number c-Myc positive cells by lesion area (million pixels) in CMTB mice compared to control. **G)** Correlation between number of c-Myc-expressing cells in a lesion and area of the lesion, in CMTB mice and in control mice. Data were analyzed using a two-tailed Student’s t-test.

Our analysis demonstrated a significant increase in c-Myc transcriptional programs in monocytes and macrophages during CMTB in mice (Figure 5C). In sum, single-cell transcriptomic analysis of myeloid cells from the CMTB model revealed elevated expression of c-Myc-associated transcriptional programs in monocytes and macrophages, linking c-Myc activity to mycobacterial persistence in vivo.

### c-Myc is associated with increased lesion size in the standard lung infection model in mice

To examine c-Myc expression in lung tissue, we conducted immunohistochemical staining with anti-c-Myc antibodies on archived lung tissue sections from prior mouse infection studies (Figure 5D). These studies compared MTB reinfection in mice with CMTB, and control mice not pre-exposed to MTB from our previously published experiments^9^. Automated image analysis identified c-Myc positive cells and normalized this count to lesion size. At 3 months post-infection, a significant increase in c-Myc-positive cell density was observed within the lesions. Further analysis revealed a positive correlation between lesion size and the number of c-Myc-positive cells. Additionally, a significant interaction was detected between lesion size and experimental group (control vs. CMTB), suggesting that the relationship between lesion size and c-Myc-positive cell count varies by group (Figure 5E). Specifically, as lesion size increases, the control group showed a greater increase in c-Myc-positive cells than the CMTB group (Figure 5F). This indicates that prior MTB exposure in the CMTB model may reduce the accumulation of c-Myc-positive cells or reduce the expression of c-Myc in cells already present as lesions enlarge, potentially contributing to reduced lesion growth. However, while this interpretation aligns with our data, causality cannot be confirmed by these findings alone. In sum, immunohistochemical analysis of lung tissue revealed that c-Myc expression positively correlates with lesion size during standard MTB lung infection model.

### c-Myc expression is spatially enriched in the immune-privileged core of human tuberculosis granulomas

A key limitation of the mouse model is the absence of granulomas, a hallmark of mycobacterial infections in humans. Granulomas with central necrosis are a defining feature of mycobacterial infections in humans and are structured to contain the pathogen^3^. These granulomas generally have a necrotic core surrounded by an outer layer of immune cells, with the inner granuloma known to be an immune-privileged site. We therefore hypothesized that c-Myc expression may be elevated in the inner granuloma compared to surrounding tissues in humans.

To test this, we analyzed c-Myc expression in human granulomas by comparing its levels relative to the necrotic center. Tissue samples were collected from seven patients diagnosed with active tuberculosis during routine clinical procedures at the University Hospital Zurich (Supplementary Table 1). We used a semi-automated tissue analysis, in which inner and outer granulomas were manually segmented followed by automatic nuclei detection and classification of mean c-Myc staining intensity (Figure 6A). We found that the inner region of the granuloma contained a significantly higher proportion - up to 30% more - of c-Myc-expressing cells compared to the outer region (Figure 6B). Moreover, the frequency of c-Myc-positive cells declined with increasing distance from the granuloma core (Figure 6C), and among those positive cells, the intensity of c-Myc expression also progressively decreased with distance. In summary, c-Myc expression is spatially associated with the immune-privileged environment of the human granuloma.

**Figure 6:**
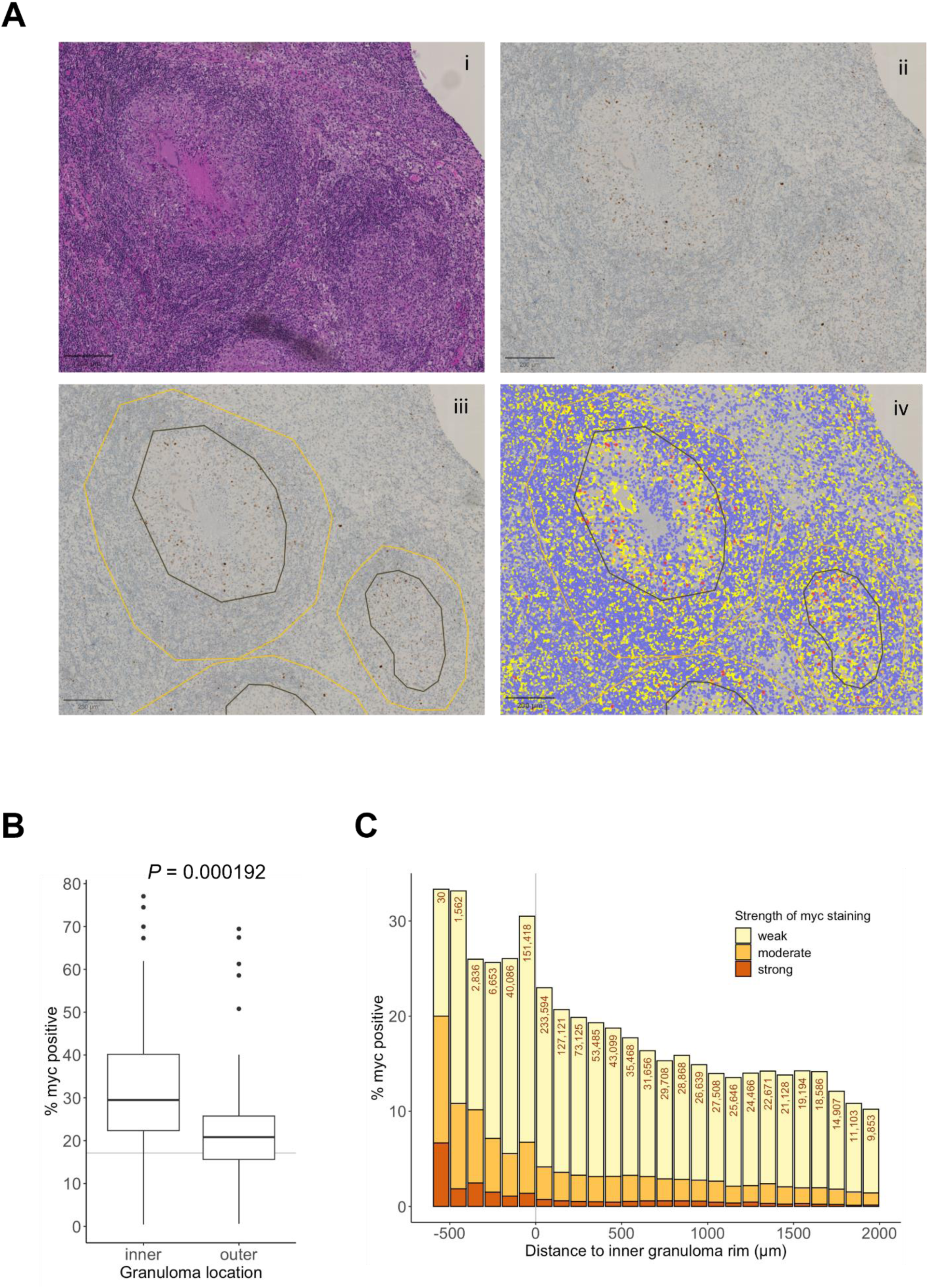
c-Myc expression is associated with the immune privileged niche in the human granuloma during active TB. **A)** Immunohistochemical stain for c-Myc (i-ii) in inner or outer granuloma. Inner and outer granulomas were manually segmented (iii) followed by automatic nuclei detection and classification of mean c-Myc staining intensity into negative (blue) or positive: weak (yellow), moderate (orange), or strong (red) (iv). **B)** Boxplot showing percent of c-Myc -positive nuclei in granuloma regions; inner granulomas had a higher mean percentage of nuclei positive for c-Myc staining (32.3%) compared to outer granulomas (23.1%). The horizontal grey line shows the percentage of positive nuclei outside granulomas (17.1%). **C)** Individual nuclei were grouped by distance to the rim of the nearest inner granuloma into 100 µm bins and the percentages of nuclei with positive staining by intensity were calculated; negative distances indicate the nucleus is within the inner granuloma. The total number of nuclei within each bin is represented at the top. Data were analyzed using a two-tailed Student’s t-test.

## Discussion

MTB remains a major cause of mortality worldwide. However, over 90% of exposed individuals do not develop active disease, indicating a significant variability in clinical outcomes likely influenced by host immune factors^29^. Based on prior findings and clinical observations, we hypothesized that the activation state of macrophages, particularly the sequence of activation relative to infection, may play a critical role in determining the outcome of MTB exposure. To test this hypothesis, we employed a controlled in vitro model to examine the effect of IFN-γ activation timing on macrophage responses to infection. Using an unbiased systems approach, we identified c-Myc as a crucial regulator of antimycobacterial activity in vitro. Inhibiting c-Myc induced a proinflammatory macrophage phenotype with enhanced antimycobacterial properties. Mechanistically, c-Myc inhibition shifted macrophage metabolism including increased mTORC1 activation, which improved MTB control.

In line with these findings, inflammation has been linked to suppression of c-Myc associated gene programs^30^. Also, very recent data suggest that overexpression of c-Myc leads to a state of chronic oxidative stress for the macrophage^31^. c-Myc is a crucial regulator in cancer and stem cell biology, known for its role in controlling cell proliferation, growth, and metabolism^32–35^. c-Myc has multiple functions in different cells populations^36–38^. In macrophages, c-Myc has been implicated in the regulation of macrophage polarization. While c-Myc has been associated with the outdated M2 macrophage polarization concept as a potential marker of M2 macrophages^39–41^, its specific functions in mature macrophages have not been thoroughly explored.

We tackled the experimental challenge of c-Myc inhibition by employing a tetracycline-inducible Omomyc system, which enabled us to differentiate macrophages into their mature form without disruption and subsequently inhibit c-Myc activity post-maturation. Omomyc, a synthetic dominant-negative regulator of c-Myc, competes with c-Myc to form dimers with Max and binds to c-Myc DNA target sequences (E-boxes) preventing the transactivation of the target genes^22^. This approach has several advantages over chemical inhibitors. It removes the need for solvents and avoids delivery challenges, as the inhibitor is produced directly inside the target cells.

Our data indicate that c-Myc signaling in mature macrophages significantly contributes to macrophage functionality even in the fully matured cell. Surprisingly, the primary differences between MTB-permissive and MTB-controlling macrophages occur not only in proinflammatory “effector” pathways but also in canonical cell cycle pathways. This finding challenges traditional views of c-Myc being key primarily for the development of cells but positions it also a key regulator of mature macrophage responses^30,42–44^ Inhibition of c-Myc does not immediately compromise cell viability but instead induces a TNF-α, iNOS, and mTORC1-driven transcriptional program known to be interconnected and to enhance macrophage function during mycobacterial infection^45^.

Alveolar macrophages, the cells first to be infected with MTB, and BMDMs represent distinct macrophage populations that differ in ontogeny, tissue localization, and baseline activation states. AMs are tissue-resident macrophages originating from fetal precursors and adapted to the lung environment, whereas BMDMs are derived from circulating monocytes and typically generated ex vivo under defined cytokine conditions^46,47^. Despite these differences, comparative analysis of transcriptional responses across these cell types can reveal conserved mechanisms of host defense. Complementing our findings in BMDMs, a review of previously published RNA-seq data from AMs during MTB infection^9^ indicated that AMs with increased bactericidal activity downregulate c-Myc-associated gene programs^28^. Notably, the transcriptional signature of these more effective AMs closely mirrors the profile observed in MTB-controlling BMDMs, including suppression of c-Myc target genes and engagement of inflammatory and metabolic pathways. This convergence suggests that c-Myc suppression may represent a generalizable feature of macrophage-mediated control of MTB, operating across ontogenically and functionally distinct macrophage subsets both in vitro and in vivo.

We found that c-Myc expression and its related transcriptional profiles are strongly associated with MTB infection and active tuberculosis in two different mouse models of MTB infection as well as in human samples from patients with confirmed active TB. Importantly, the spatial pattern of c-Myc - with up to 30% of cells in the inner granuloma expressing c-Myc during active TB - aligns with the well-established structure of granulomas, where the core is myeloid cell rich and immune-privileged^3^. Since c-Myc is recognized in cancer research as a key regulator of immune privilege, this observation appears intuitive, as granulomas are locally immune-suppressed regions^34^. Additionally, previous studies have identified c-Myc as an essential factor in the formation of multinucleated giant cells, another feature of mycobacterial infections^48^. While we did not specifically examine multinucleated giant cells in this study, and instead focused on c-Myc expression per nucleus rather than per cell, our findings suggest that c-Myc could be a critical mediator of mycobacterial persistence in the host.

The crucial role of c-Myc in survival and inflammatory response makes it a target for specialized intracellular pathogens that must prevent pro-inflammatory activation and promote survival of the infected cells in order to proliferate. Indeed, upregulation of c-Myc has been reported in cells infected with *Chlamydia trachomatis*^49–51^, *Salmonella typhimurium*^52^ and the protozoan pathogen *Leishmania donovani*^53^, demonstrating that this mechanism evolved in different intracellular pathogens. Furthermore, repression of c-Myc signaling alone has been proved to reduce the growth of *Chlamydia*^50,51^ and *Leishmania*^53^. These results suggest that c-Myc may play a significant role in intracellular infections in general.

Our data indicate that c-Myc inhibition leads to upregulation of several metabolic pathways including mTORC1. Given that mTORC1 directly and indirectly regulates numerous cellular processes^45^ - including iNOS and TNF-α expression, metabolic reprogramming, autophagy, and trained immunity - it is not surprising that reduced mTORC1 activity diminishes antimycobacterial responses. This partial reversal of the enhanced effects observed with c-Myc inhibition suggests that mTORC1 activity is a critical mediator of the functional gain induced by c-Myc suppression. The metabolic profile resembles the trained immunity phenotype, which likewise leads to a gain of antimicrobial functions in myeloid cells^54^. Interestingly, we did not observe a canonical switch towards glycolysis, which is typically considered a hallmark of activated, pro-inflammatory macrophages with enhanced antimicrobial activity^45,55,56^. Together these findings suggest that metabolic adaptations may influence the course of MTB infections, emphasizing the importance of metabolic reprogramming for defense. They also highlight the interaction between c-Myc and mTORC1, which could enhance the function of macrophages. Furthermore, the presented results provide a potential explanation for the observation that despite the presence of IFN-γ or IFN-γ-producing cells at the site of infection ^15,16^, active disease can occur. Being that higher expression of c-Myc – as seen in granulomas - limits the macrophages ability to respond properly to the pathogen.

The primary limitation of this study is the absence of in vivo data to demonstrate the effects of c-Myc perturbation on MTB infection, such as, for example, in a mouse model. Although future studies will address this, we have observed strong in vivo associations, including the immune privileged core of human granulomas, where MTB is located^3,8,17^.

In conclusion, our data show that c-Myc signaling plays a major role in influencing antimycobacterial responses in macrophages in vitro. The strong in vivo association in both small animal models and in human tissues from patients suffering from active TB support further investigation into the interactions between c-Myc and mycobacterial infections.

## Materials

Key Resource Table

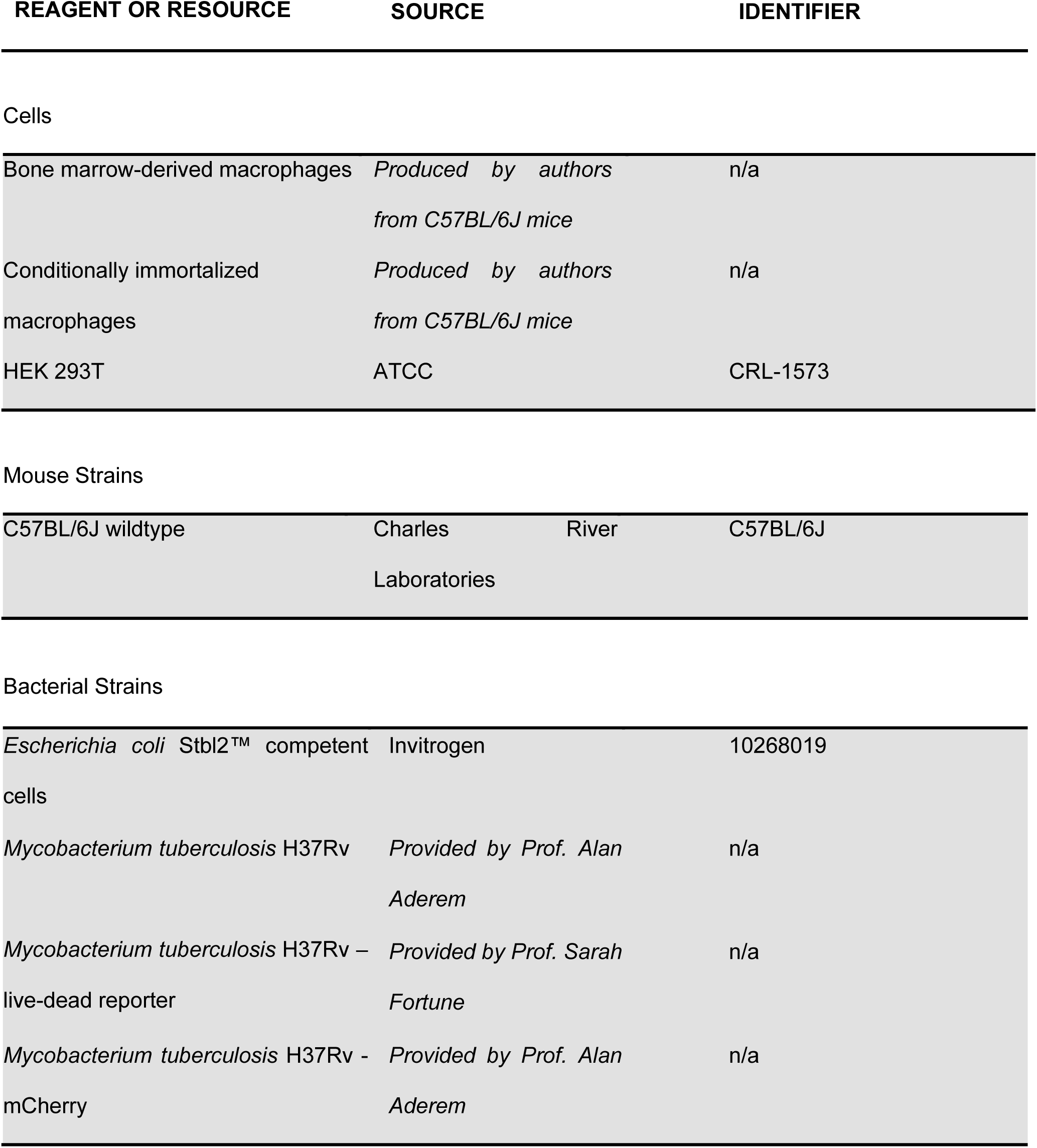
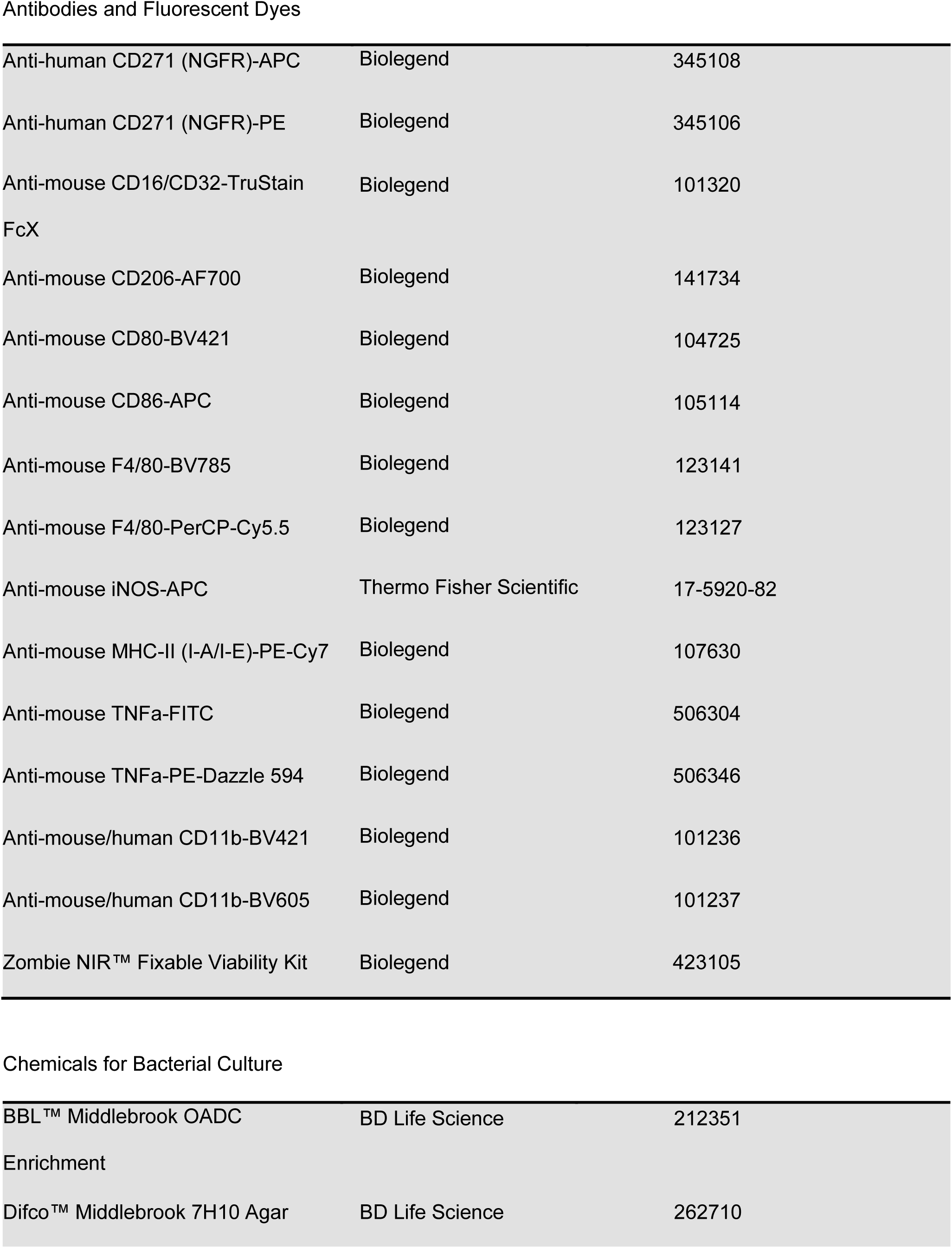
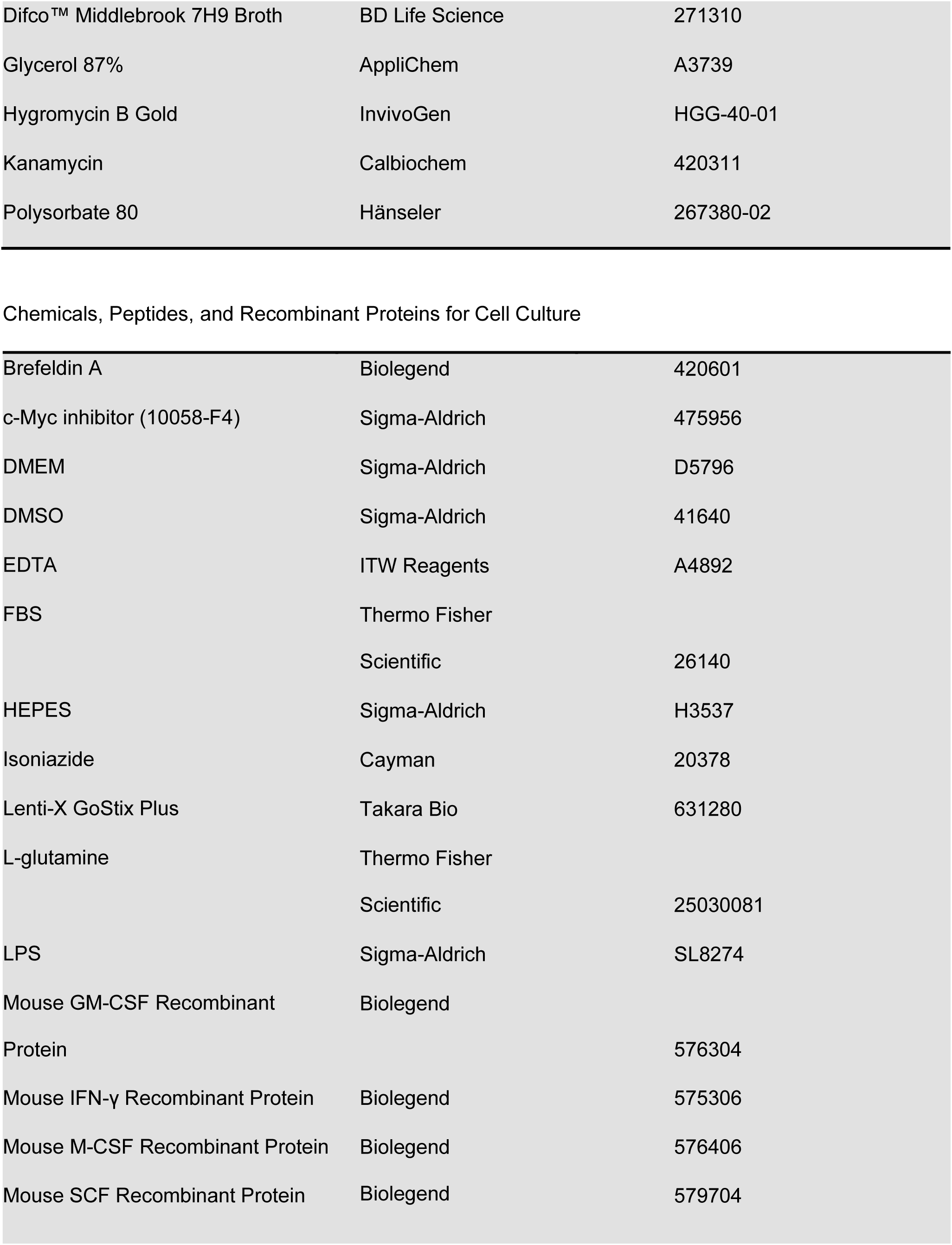
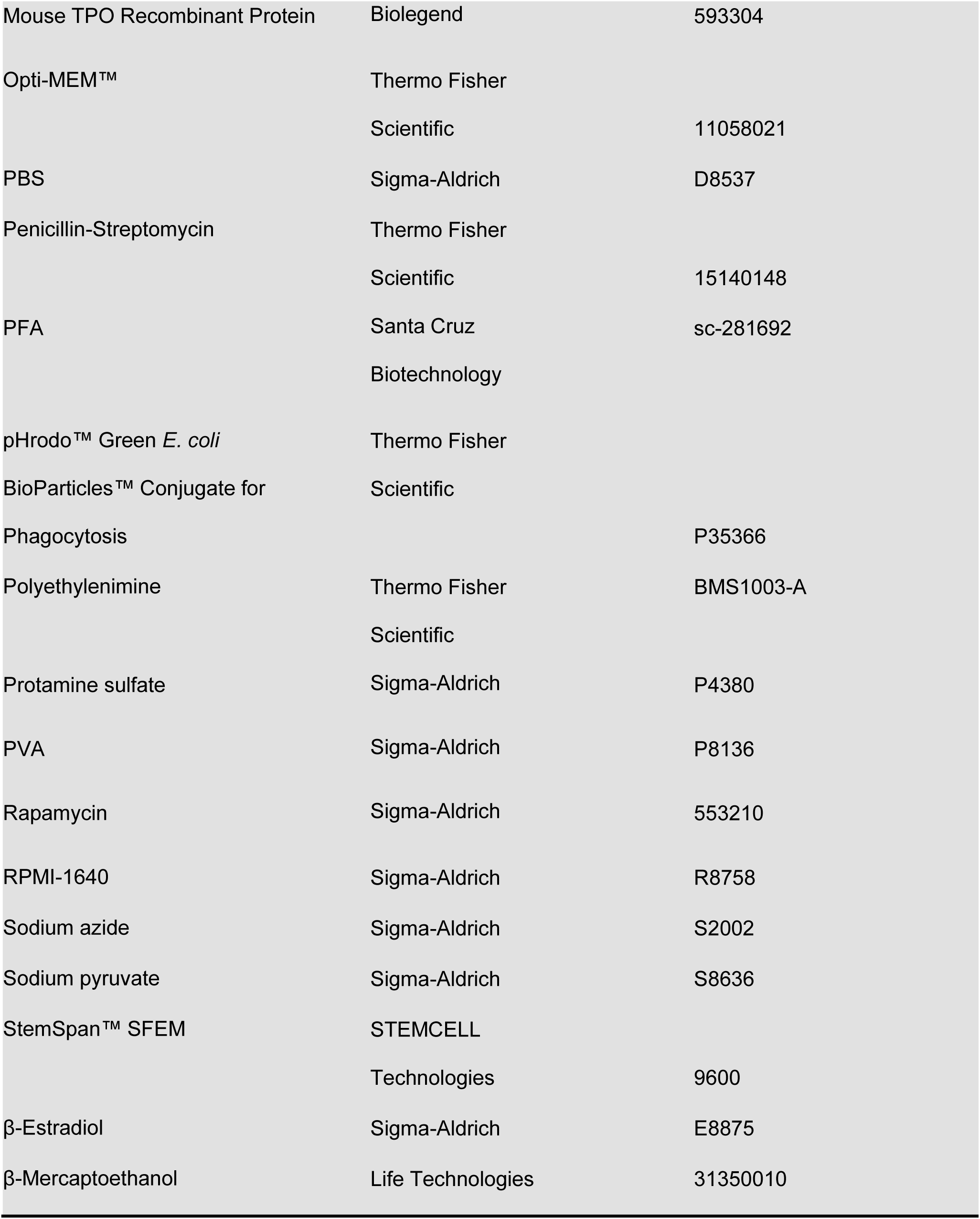
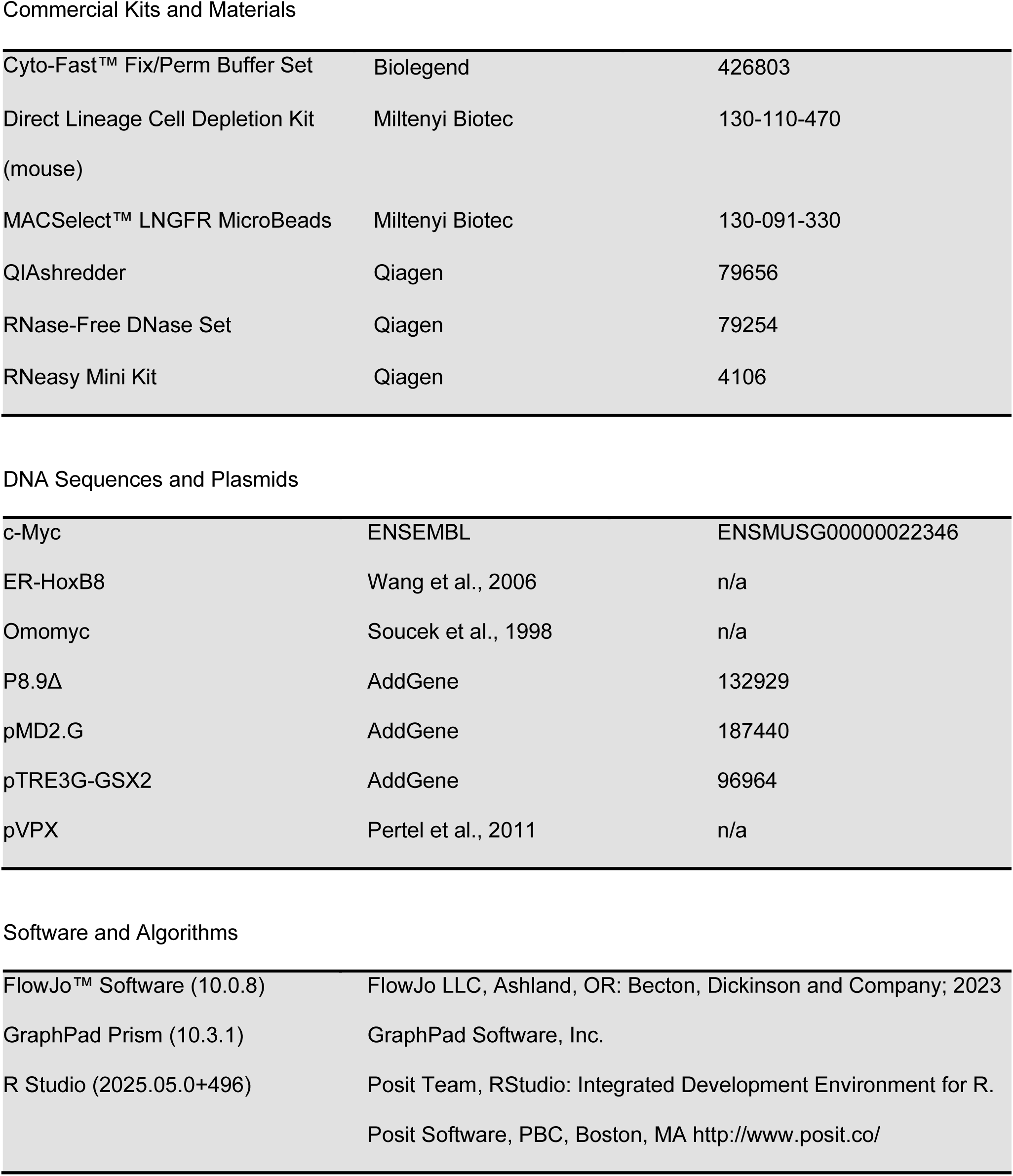

| REAGENT OR RESOURCE | SOURCE | IDENTIFIER |
| --- | --- | --- |
| Cells |  |  |
| Bone marrow-derived macrophages | *Produced by authors from C57BL/6J mice* | n/a |
| Conditionally immortalized macrophages | *Produced by authors from C57BL/6J mice* | n/a |
| HEK 293T | ATCC | CRL-1573 |
| Mouse Strains |  |  |
| C57BL/6J wildtype | Charles River Laboratories | C57BL/6J |
| Bacterial Strains |  |  |
| *Escherichia coli* Stbl2™ competent cells | Invitrogen | 10268019 |
| *Mycobacterium tuberculosis* H37Rv | *Provided by Prof. Alan Aderem* | n/a |
| *Mycobacterium tuberculosis* H37Rv – live-dead reporter | *Provided by Prof. Sarah Fortune* | n/a |
| *Mycobacterium tuberculosis* H37Rv - mCherry | *Provided by Prof. Alan Aderem* | n/a |
| Antibodies and Fluorescent Dyes |  |  |
| Anti-human CD271 (NGFR)-APC | Biolegend | 345108 |
| Anti-human CD271 (NGFR)-PE | Biolegend | 345106 |
| Anti-mouse CD16/CD32-TruStain FcX | Biolegend | 101320 |
| Anti-mouse CD206-AF700 | Biolegend | 141734 |
| Anti-mouse CD80-BV421 | Biolegend | 104725 |
| Anti-mouse CD86-APC | Biolegend | 105114 |
| Anti-mouse F4/80-BV785 | Biolegend | 123141 |
| Anti-mouse F4/80-PerCP-Cy5.5 | Biolegend | 123127 |
| Anti-mouse iNOS-APC | Thermo Fisher Scientific | 17-5920-82 |
| Anti-mouse MHC-II (I-A/I-E)-PE-Cy7 | Biolegend | 107630 |
| Anti-mouse TNFa-FITC | Biolegend | 506304 |
| Anti-mouse TNFa-PE-Dazzle 594 | Biolegend | 506346 |
| Anti-mouse/human CD11b-BV421 | Biolegend | 101236 |
| Anti-mouse/human CD11b-BV605 | Biolegend | 101237 |
| Zombie NIR™ Fixable Viability Kit | Biolegend | 423105 |
| Chemicals for Bacterial Culture |  |  |
| BBL™ Middlebrook OADC Enrichment | BD Life Science | 212351 |
| Difco™ Middlebrook 7H10 Agar | BD Life Science | 262710 |
| Difco™ Middlebrook 7H9 Broth | BD Life Science | 271310 |
| Glycerol 87% | AppliChem | A3739 |
| Hygromycin B Gold | InvivoGen | HGG-40-01 |
| Kanamycin | Calbiochem | 420311 |
| Polysorbate 80 | Hänseler | 267380-02 |
| Chemicals, Peptides, and Recombinant Proteins for Cell Culture |  |  |
| Brefeldin A | Biolegend | 420601 |
| c-Myc inhibitor (10058-F4) | Sigma-Aldrich | 475956 |
| DMEM | Sigma-Aldrich | D5796 |
| DMSO | Sigma-Aldrich | 41640 |
| EDTA | ITW Reagents | A4892 |
| FBS | Thermo Fisher |  |
|  | Scientific | 26140 |
| HEPES | Sigma-Aldrich | H3537 |
| Isoniazide | Cayman | 20378 |
| Lenti-X GoStix Plus | Takara Bio | 631280 |
| L-glutamine | Thermo Fisher |  |
|  | Scientific | 25030081 |
| LPS | Sigma-Aldrich | SL8274 |
| Mouse GM-CSF Recombinant | Biolegend |  |
| Protein |  | 576304 |
| Mouse IFN-γ Recombinant Protein | Biolegend | 575306 |
| Mouse M-CSF Recombinant Protein | Biolegend | 576406 |
| Mouse SCF Recombinant Protein | Biolegend | 579704 |
| Mouse TPO Recombinant Protein | Biolegend | 593304 |
| Opti-MEM™ | Thermo Fisher |  |
|  | Scientific | 11058021 |
| PBS | Sigma-Aldrich | D8537 |
| Penicillin-Streptomycin | Thermo Fisher |  |
|  | Scientific | 15140148 |
| PFA | Santa Cruz | sc-281692 |
|  | Biotechnology |  |
| pHrodo™ Green *E. coli* | Thermo Fisher |  |
| BioParticles™ Conjugate for | Scientific |  |
| Phagocytosis |  | P35366 |
| Polyethylenimine | Thermo Fisher | BMS1003-A |
|  | Scientific |  |
| Protamine sulfate | Sigma-Aldrich | P4380 |
| PVA | Sigma-Aldrich | P8136 |
| Rapamycin | Sigma-Aldrich | 553210 |
| RPMI-1640 | Sigma-Aldrich | R8758 |
| Sodium azide | Sigma-Aldrich | S2002 |
| Sodium pyruvate | Sigma-Aldrich | S8636 |
| StemSpan™ SFEM | STEMCELL |  |
|  | Technologies | 9600 |
| β-Estradiol | Sigma-Aldrich | E8875 |
| β-Mercaptoethanol | Life Technologies | 31350010 |
| Commercial Kits and Materials |  |  |
| Cyto-Fast™ Fix/Perm Buffer Set | Biolegend | 426803 |
| Direct Lineage Cell Depletion Kit<br>(mouse) | Miltenyi Biotec | 130-110-470 |
| MACSelect™ LNGFR MicroBeads | Miltenyi Biotec | 130-091-330 |
| QIAshredder | Qiagen | 79656 |
| RNase-Free DNase Set | Qiagen | 79254 |
| RNeasy Mini Kit | Qiagen | 4106 |
| DNA Sequences and Plasmids |  |  |
| c-Myc | ENSEMBL | ENSMUSG00000022346 |
| ER-HoxB8 | Wang et al., 2006 | n/a |
| Omomyc | Soucek et al., 1998 | n/a |
| P8.9Δ | AddGene | 132929 |
| pMD2.G | AddGene | 187440 |
| pTRE3G-GSX2 | AddGene | 96964 |
| pVPX | Pertel et al., 2011 | n/a |

Software and Algorithms
|  |  |
| --- | --- |
| FlowJo™ Software (10.0.8) | FlowJo LLC, Ashland, OR: Becton, Dickinson and Company; 2023 |
| GraphPad Prism (10.3.1) | GraphPad Software, Inc. |
| R Studio (2025.05.0+496) | Posit Team, RStudio: Integrated Development Environment for R.<br>Posit Software, PBC, Boston, MA http://www.posit.co/ |

### Experimental models

#### Mycobacterium tuberculosis strains

MTB strain H37Rv and H37Rv mCherry fluorescent MTB strains were kindly provided by Prof. Alan Aderem. MTB H37Rv live-dead reporter strain was provided by Prof. Sarah Fortune. All strains were cultured in Difco Middlebrook 7H9 supplemented with 10%(v/v) BBL Middlebrook OADC and polysorbate 80 in roller bottles or Difco Middlebrook 7H10 Agar supplemented with 10%(v/v) BBL Middlebrook OADC and 0.5% (v/v) Glycerol 87%. Kanamycin (30 μg/mL) and hygromycin B (50 μg/mL) were added to the MTB H37Rv mCherry strain and the MTB H37Rv live-dead strain respectively. Bacteria were quantified by spectrophotometry (OD 600nm). To assess transcriptional activity of the MTB H37Rv live-dead strain^20^ (after macrophage infection), doxycycline (1 µg/mL) was added to the culture medium for 24 hours.

#### Mouse strains

C57BL/6J female mice were bought from Charles River Laboratories and were used as donor for bone marrow macrophages production. All procedures were approved by the cantonal animal experimentation authorities and followed international guidelines for laboratory animal experimentation (Canton of Zurich, licenses ZH138/19 and ZH041/21). All mice were maintained on a 12-hour light-dark cycle under clean, conventional conditions and with free access to water and food.

## Methods

### Cell culture and cell isolation from bone marrow

HEK 293T cells were cultured in Dulbecco’s Modified Eagle Medium (DMEM) supplemented with 10% fetal bovine serum (FBS), 2mM L-glutamine, and 1% Penicillin-Streptomycin (10’000U/mL; P/S). Bone marrow cells were harvested as described by Amend et al.^57^. To induce differentiation into BMDMs, bone marrow cells were cultured in BMDM differentiation medium, consisting of Roswell Park Memorial Institute (RPMI) 1640 medium supplemented with 10% FBS, 2mM L-glutamine, 1% P/S, and 50 ng/mL macrophage colony-stimulating factor (M-CSF), at a density of 2×10^6^ cells/mL. The medium was refreshed on day 3. On day 6, non-adherent suspension cells were discarded, and adherent cells were detached (with cold PBS + 5 mM EDTA by pipetting) for downstream experiments. To isolate hematopoietic stem cells (HSCs) and immune progenitors from the bone marrow, negative magnetic selection was performed using the Direct Lineage Cell Depletion Kit following the manufacturer’s protocol. Bone marrow hematopoietic progenitors were grown in 6-well plates with HSC medium, consisting of StemSpan SFEM medium supplemented with 1 mg/mL polyvinyl alcohol (PVA), 50 ng/mL thrombopoietin (TPO), and 100 ng/mL stem cell factor (SCF), adapted from Wilkinson et al^58^. For one proof of concept experiment, lineage depleted cells were immortalized using an estrogen regulated HoxB8 (ER-HoxB8) lentiviral vector as described by Wang et al.^59^. Those cells can be kept in a monocyte precursor stage for over 850 generations in immortalization medium consisting of RPMI 1640 medium containing 10% FBS, 2mM L-glutamine, 1% P/S, 10 mM HEPES, 50 µM β-mercaptoethanol, 1 mM sodium pyruvate, 2 µM β-estradiol, and 20 ng/mL granulocyte-macrophage colony-stimulating factor (GM-CSF). Differentiation into mature macrophages occurs within 4 to 6 days in complete differentiation medium.

### Cloning of the Tet-on system into lentiviral vectors

rtTA V16, an improved version of the reverse tetracycline transactivator used in Tet-on systems was developed by Das et al.^60^. TRE3G inducible promoter was obtained from plasmid pTRE3G-GSX2. Omomyc peptide sequence was obtained from Soucek et al.^61^, and inserted in the construct. The coding sequence of the murine version of c-Myc was obtained from the ENSEMBL database.

### Production of lentiviral particles

Lentiviruses were generated by transfecting HEK 293T cells with lentiviral vectors in Opti-MEM using polyethylenimine (PEI) as the transfection reagent. The transfection mix included p8.9Δ (containing the coding sequences for gag/pol, rev, and tat under the control of the immediate/early promoter enhancer of cytomegalovirus), pMD2.G (encoding vesicular stomatitis virus glycoprotein), and a VPX-expressing plasmid (as described by Pertel et al.^62^) in a 1:3 DNA-to-PEI ratio. The medium was replaced 16 hours post-transfection with RPMI supplemented with 5% FBS. Supernatants containing lentiviral particles were collected and filtered 48 hours after transfection. Lentiviral production was confirmed using the Lenti-X GoStix Plus assay. Viral particles were subsequently concentrated by centrifugation at 20’000 g for 2 hours, resuspended in PBS, and stored at −80°C until further use.

### Transduction of mouse HSCs and myeloid progenitors

HSCs and immune progenitors were isolated from mice as described above and cultured overnight in HSC medium (described above). For transduction, the medium was prepared by supplementing the HSC medium with 10 µg/mL protamine and the appropriate lentiviruses at a multiplicity of infection (MOI) of 10. Cells were resuspended in the transduction medium at a density of 1×10^6^ cells/mL. After 12 hours of incubation, the cells were pelleted by centrifugation, washed twice with PBS, and resuspended in BMDM differentiation medium to promote differentiation into macrophages. After 5-6 days of differentiation, the transduced cells expressing the nerve growth factor receptor (NGFR) were magnetically enriched using MACSelect LNGFR MicroBeads according to the manufacturer’s instructions. NGFR⁺ cells were counted and used for downstream experiments.

### Macrophage infection assays

BMDMs were seeded in differentiation medium in 96-well plates, 5x10^4^ cells per well. After 24 hours, the cells were treated with IFN-γ 25ng/mL according to conditions. On the day of infection, bacteria in log-phase (OD 0.1-0.5) were pelleted by centrifugation and resuspended in RPMI and used to infect BMDMs. BMDMs were infected at a MOI of 5 bacilli per macrophage for 1 hour at 37°C and washed twice with PBS to remove extracellular bacteria. The cells were provided fresh BMDM differentiation medium and with or without IFN-γ according to conditions. For assays without IFN-γ but involving Tet-on constructs (c-Myc- and Omomyc-expressing BMDMs), the differentiation medium was supplemented with doxycycline (100 ng/mL) 24 hours prior to infection and maintained throughout the experiment. For rapamycin experiments, cells were pretreated for 30 minutes in BMDM differentiation medium supplemented with or without 10 µM rapamycin, depending on the experimental condition. All conditions received the same concentration of DMSO (0.1%). Doxycycline (final concentration 100 ng/mL) was then added where indicated. Rapamycin treatment was also maintained throughout the experiment. For all experiments medium was changed latest every second day according to the conditions. At day 0 (1 hour after infection), 3 and 5 the macrophages were detached from the wells with cold PBS + 5 mM EDTA by pipetting gently. Half of the suspension with the cells were either used for flow cytometry staining or lysed with 0.05% SDS in water, and the lysate was plated on 7H10 agar plates in serial dilutions. After 3 weeks of incubation at 37°C colonies were counted to measure the anti-bacterial activity of macrophages.

### Flow cytometry staining

Staining was performed in FACS buffer (PBS + 2 mM EDTA + 2% FBS + 0.1% sodium azide) containing CD16/CD32 blocker (FcX), Zombie NIR dye and the fluorescent antibodies (= master mix). Suspension cells were pelleted by centrifugation at 300 g for 5 minutes and washed once prior to staining. BMDMs were detached by removing the media, washing with PBS, and incubating the cells for 5-10 minutes in cold PBS + 5 mM EDTA, followed by gentle pipetting to detach the cells. Cells were incubated in the master mix for 20 minutes in the fridge. Cells were washed and fixed in 2–4% paraformaldehyde (PFA) at room temperature for 20 minutes, with particular emphasis on MTB-infected cells. For intracellular staining, the Cyto-Fast Fix/Perm Buffer Set was used according to the manufacturer’s instructions before fixation. When stimulation was performed, cells were stimulated for 6 hours with 10 mg/mL lipopolysaccharide (LPS) and 1X Brefeldin A (BFA) before being stained. Acquisition was conducted either at a CytoFLEX S (Beckman Coulter) or LSR Fortessa (BD Biosciences).

### Assay with heat-killed MTB and pHrodo Green

BMDMs differentiated and transduced as described above were incubated with heat-killed MTB at a ratio of 1:2 (BMDM:bacteria) for 1 hour at 37°C. Heat inactivation was performed by incubating mCherry-expressing MTB H37Rv cultures at 80°C for 20 minutes using a thermoblock. After incubation, the cells were washed twice with PBS and fresh BMDM differentiation medium was added. After 24 hours, the medium was replaced with 100 µL of fresh differentiation medium. pHrodo Green *E. coli* bioparticles (pre-aliquoted at 1 mg/mL) were added at 5 µL per well. Cells were incubated for 1 hour at 37°C in an incubator without CO_2_, as CO_2_ can alter the pH of the medium and interfere with pHrodo fluorescence. After incubation, the cells were washed with PBS and resuspended in FACS buffer. Samples were immediately acquired by flow cytometry without fixation and pHrodo fluorescence was detected in the GFP channel.

### Mouse infection experiments

All mouse experiments regarding the CMTB model have been previously reported in Nemeth *et al.*^9^. For completeness, we briefly describe the establishment of CMTB using intradermal ear injection and aerosol infection. Intradermal infections involved administering 10’000 CFU of logarithmic-phase MTB H37Rv in 10 μL PBS using a 10 μL Hamilton syringe into mice anesthetized with ketamine, following a modified protocol. Aerosol infections were performed using a diluted frozen stock of kanamycin-resistant MTB H37Rv in an aerosol infection chamber (Glas-Col). Bacterial load was assessed by plating serial dilutions of homogenized tissue on kanamycin-containing and antibiotic-free plates to differentiate between the sources of infection.

### RNA sequencing

For bulk RNA-seq experiments, 1×10⁶ BMDMs that had differentiated for 6 days were seeded in 12-well plates. Depending on the experimental conditions, addition of IFN-γ or doxycycline, as well as in vitro infections, were performed as described. For RNA isolation, samples were collected 6 and/or 24 hours post-infection or post-induction. RNA was isolated using the RNeasy Mini Kit, in combination with QIAshredder homogenization columns and the RNase-Free DNase Set, according to the manufacturer’s instructions. RNA quality was assessed using a Fragment Analyzer (Agilent), and concentrations were measured using a Qubit (1.0) Fluorometer (Life Technologies). Only samples with 260/280 ratios between 1.8 and 2.1 and 28S/18S ratios of 1.5–2 were used for downstream processing.

Library construction, sequencing, and parts of the data analysis were performed by the Functional Genomics Center Zurich. mRNA libraries were prepared using the TruSeq Stranded mRNA Library Prep Kit (Illumina), which includes poly-A selection, reverse transcription, fragmentation, end-repair, and adapter ligation. Fragments with adapters on both ends were enriched via PCR using unique dual indexing (UDI). Library quality and concentration were validated using a Qubit (1.0) Fluorometer (Life Technologies) and Fragment Analyzer (Agilent). Libraries were normalized to 10 nM in 10 mM Tris-Cl (pH 8.5) with 0.1% Tween 20. Sequencing was performed as single-end 100 bp reads on the Illumina NovaSeq 6000 platform.

Raw reads were quality filtered and trimmed using fastp. Transcript quantification was performed by pseudo-aligning reads to the mouse genome (GRCm38.p6 with GENCODE release M23) using Kallisto. Differential expression analysis was performed in R using “DESeq2”^63^. PCA, heatmaps, and other visualizations were generated in RStudio using packages “ComplexHeatmap”^64^, “circlize”^65^, “EnhancedVolcano”, “patchwork”, and “tidyverse”^66^. GSEA was conducted with “fgsea”^67^ using the Mouse Hallmark gene set from MSigDB^68–70^ (v2024.1.Mm) and pathway-gene annotations were supplied by “org.Mm.eg.db”.

### Single cell RNA sequencing

Single cell RNA sequencing was performed by our collaborators in the United States. Lymph nodes were obtained from mice with and without CMTB (n = 5 per condition). Following isolation, samples underwent methanol fixation according to protocol “CG000136 Rev. D” from 10X Genomics. Briefly, samples were filtered using 70 µm filters, and red blood cells were lysed with ACK lysis buffer. Post-lysis, cells were resuspended in 1 mL of ice-cold PBS with a wide-bore pipette, then transferred to a 1.5 mL low-bind Eppendorf tube. After centrifugation at 700 g for 5 minutes at 4°C, the supernatant was carefully discarded, and the cell pellet was washed twice with PBS, counted, and resuspended in 200 µL of ice-cold PBS per 1×10^6^ cells. To achieve an 80% final methanol concentration, 800 µL of ice-cold methanol was gradually added to the suspension. The samples were incubated at -20°C for 30 minutes and stored at -80°C for up to 6 weeks prior to rehydration.

For rehydration, samples were thawed at 4°C, pelleted at 1’000 g for 10 minutes at 4°C, and resuspended in 50 µL of a wash-resuspension buffer (0.04% BSA, 1mM DTT, 0.2U/µL Protector RNAse Inhibitor in 3X SSC buffer) to achieve a final concentration of approximately 1’000 cells/µL, accounting for an estimated 75% cell loss.

Libraries were constructed using the Next GEM Single Cell 3’ Reagent Kits v3.1 (Dual Index) from 10X Genomics, following the manufacturer’s protocol. Sequencing data were aligned to the mouse genome (mm10), and unique molecular identifier (UMI) counts were calculated using the Cell Ranger pipeline. Data processing, integration, and downstream analysis were conducted in Seurat v3, with quality control filtering to exclude droplets containing fewer than 200 detected genes, more than 4’000 detected genes (to discriminate against doublets), or more than 5% mitochondrial gene expression. Genes detected in fewer than three cells across all samples were excluded. Cell-type annotation leveraged the Immgen database as a reference, implemented with the “SingleR”^71^ package. Additionally, pseudotime trajectory analysis was performed using “SeuratWrappers” and “Monocle3”^72^. Data visualization was facilitated using the “Seurat”^73^, “tidyverse”^66^, “cowplot”, and “viridis” R packages.

### Mouse histology

Immunohistochemistry for c-Myc was performed at the University of Washington’s Histology and Imaging Core. The scanned images were exported as TIFF files and uploaded to ImageJ. Areas of consolidation were identified on images converted to 8-bit grayscale and thresholded using the “Triangle” method. After signal noise reduction with a median filter, sequential minimum and maximum filters were used to identify areas of true consolidation. c-Myc^+^ cells were identified by color-thresholding to identify brown (DAB^+^) cells against the background of blue nuclei and these were automatically counted within each consolidated lesion by using the analyze particles command. The c-Myc^+^ cell density within each consolidated lesion was calculated as the number of c-Myc^+^ cells per million pixels.

### Human histology

Patients with confirmed tuberculosis (culture- and/or PCR-positive) who had a histological specimen available and had given written consent for their routinely obtained specimens to be used for scientific research were included after ethical clearance was given by the Zurich Ethics Committee (BASEC no. 2023-00800). The specimens were stained using routine clinical protocols with hematoxylin and eosin (H&E) and an anti-c-Myc antibody. The tissues were fixed, photographed and transformed into NDPI files, which were loaded into QuPath and granulomas (both caseating and non-caseating) were identified on H&E stained slides^74^. Manual segmentation with QuPath’s polygon tool was used to create annotations for inner and outer; the inner region was subtracted from the outer region to create non-overlapping annotations. To calculate the percentage of non-epithelial cells outside granulomas that express c-Myc, an automatic threshold was applied to the c-Myc stained slides based on the average RGB intensity to identify all tissue. Areas of epithelium and the previously defined granuloma regions were excluded from the background tissue region. Additional annotations were created to exclude areas of artifact (e.g., folded tissue). Positive cell detection with the following parameters (’{“detectionImageBrightfield”:“Opticaldensitysum”,“requestedPixelSizeMicrons”:0.5,“backgroundRadiusMicr ons”:15.0,“backgroundByReconstruction”:true,“medianRadiusMicrons”:0.0,“sigmaMicrons”:1.5,“minAreaMicr ons”:10.0,“maxAreaMicrons”:400.0,“threshold”:0.12,“maxBackground”:2.0,“watershedPostProcess”:true,“ex cludeDAB”:false,“cellExpansionMicrons”:2.0,“includeNuclei”:true,“smoothBoundaries”:true,“makeMeasurem ents”:true,“thresholdCompartment”:“Nucleus: DAB OD mean”,“thresholdPositive1”:0.18,“thresholdPositive2”:0.3,“thresholdPositive3”:0.5,“singleThreshold”:false}’) was used to automatically identify nuclei and classify them into negative, positive (weak), positive (moderate), or positive (strong) c-Myc staining. Separate quantifications at the annotation (i.e. granuloma region) level and the detection (i.e. nuclei) level were exported and analyzed using R.

### Lead contact and material availability

Requests for further information and resources should be directed to and will be fulfilled by the lead contact, Johannes Nemeth. RNA data are available from NCBI Gene Expression Omnibus under accession number GSEXXXXX (tba).

## Author contributions

JN conceived the study and co-designed the experiments with ES and CD. ES, CD, RW, DM, and DR performed the experiments. KK, JHR, LB, and GSO analyzed (single-cell) RNA sequencing and histology data and provided critical insights. SB advised on plasmid design and lentivirus production. MG, PS, AHD, and RFS provided critical advice on project progression and data analysis. The manuscript was prepared by CD, JN, and ES and reviewed by all authors.

## Acknowledgements

The work was supported by the Swiss National Science Foundation (SNSF) grant 310030_200407 to JN. Work in the laboratory of PS is supported by SNSF grant 310030_197699/1, the Federal Office of Public Health (3632001500), and University of Zurich. The work of collaborators from the United States was supported by the National Institute of Allergy and Infectious Diseases of the National Institutes of Health, under awards U19AI135976 and 75N93019C00070. The authors used OpenAI’s ChatGPT to support language editing and improve the clarity of the manuscript. After using this tool, the authors reviewed and edited the content as necessary and take full responsibility for the content of the published article. Transcriptomics was performed at the Functional Genomics Centre Zurich (FGCZ) of University of Zurich and ETH Zurich.

## Supplementary Materials

**Supplementary Figure 1.**
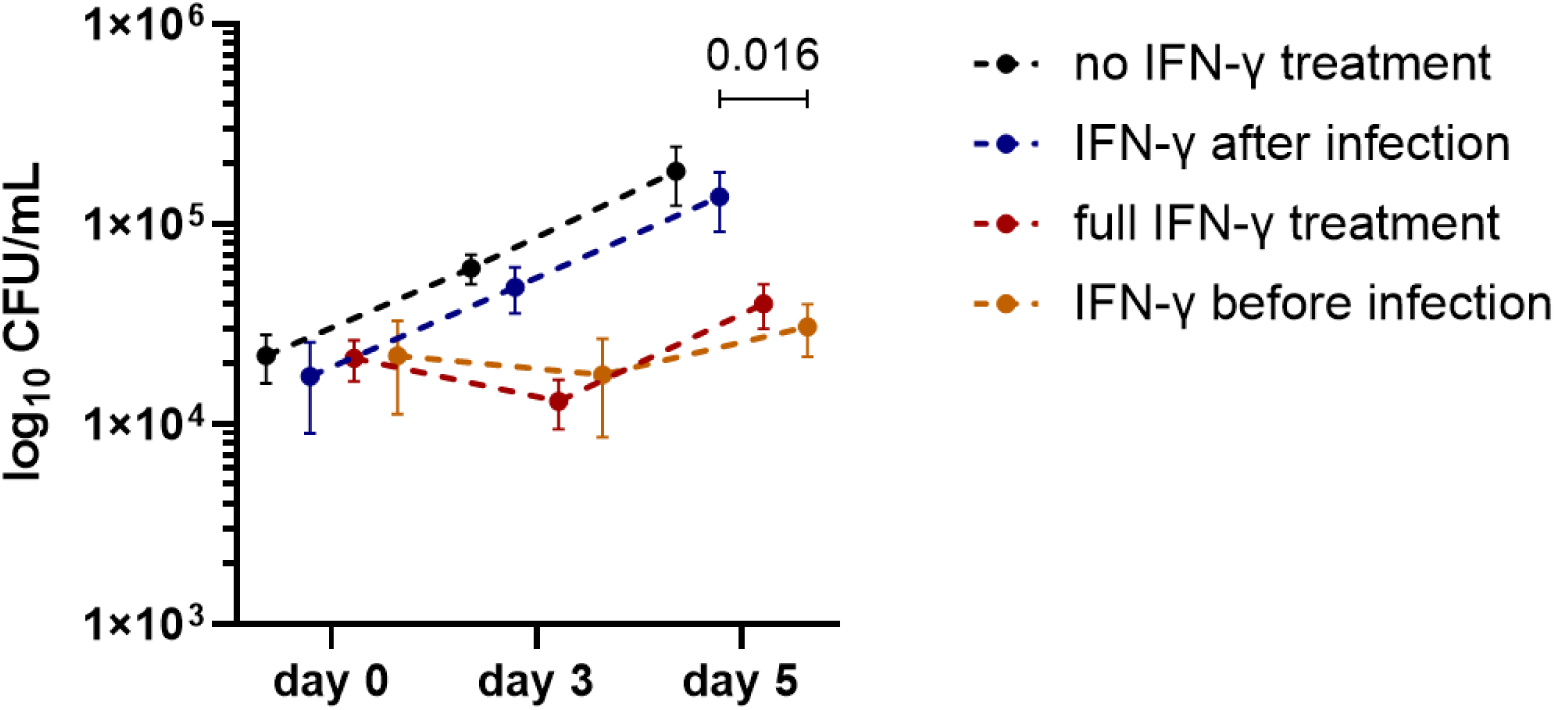
IFN-γ activation before infection improves control of MTB growth in BMDMs. Quantification in colony forming units per milliliter (CFU/mL) of intracellular MTB after IFN-γ activation at different timepoints. Measurements were taken immediately after phagocytosis (day 0) and at 3 and 5 days post-infection. Data of one experiment is shown. *P* values were determined by unpaired two-tailed Student’s t-test (before vs. after infection) and corrected for multiple testing using the two-stage linear step-up method of Benjamini–Krieger–Yekutieli.

**Supplementary Figure 2.**
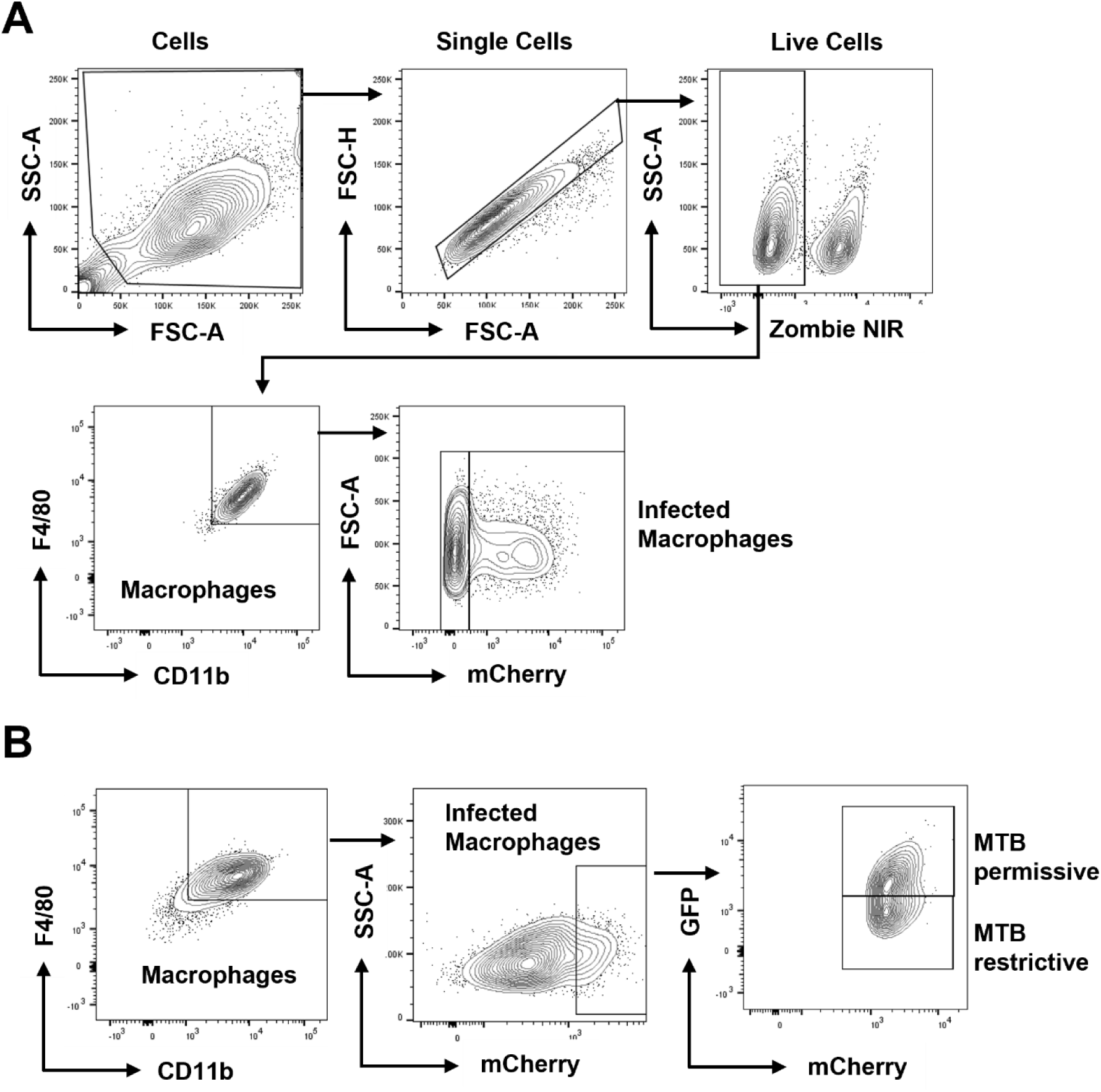
Gating strategy. Flow cytometry gating strategy of infected BMDMs infected with **A)** mCherry-expressing MTB H37Rv and **B)**MTB H37Rv live-dead reporter strain.

**Supplementary Figure 3.**
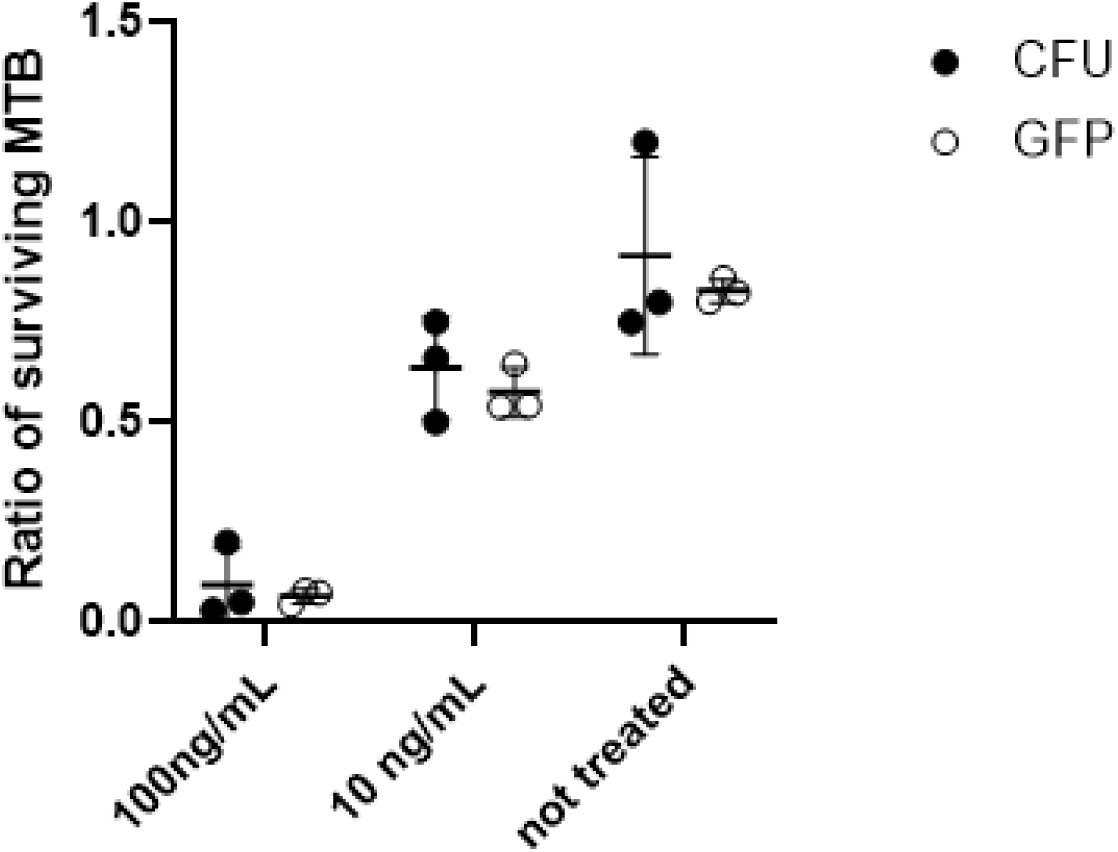
Validation of MTB survival analysis using flow cytometry and CFU assays. To validate the GFP-based flow cytometry assay against the colony forming unit (CFU) assay, we infected 50’000 BMDMs per sample with the MTB H37Rv live-dead strain (MOI 5). Cells were incubated for 24 h, followed by activation with doxycycline (1 µg/mL) for an additional 24 h. Subsequently, one half of the cell samples were lysed for CFU quantification, while the remaining cells were analyzed for GFP^+^ infected cells by flow cytometry. CFU ratios, comparing counts at baseline to those at the specified time point, were plotted to assess bacterial survival and for comparison with the MTB H37Rv live-dead strain. To establish a baseline, BMDMs infected with doxycycline-preactivated live-dead bacteria were immediately subjected to either CFU or flow cytometry analysis post-infection. MTB survival was assessed by exposing infected cells to varying concentrations of the antimycobacterial drug isoniazid (100 ng/mL, 10 ng/mL, and no antibiotic), providing a range of survival percentages for comparative analysis.

**Supplementary Figure 4.**
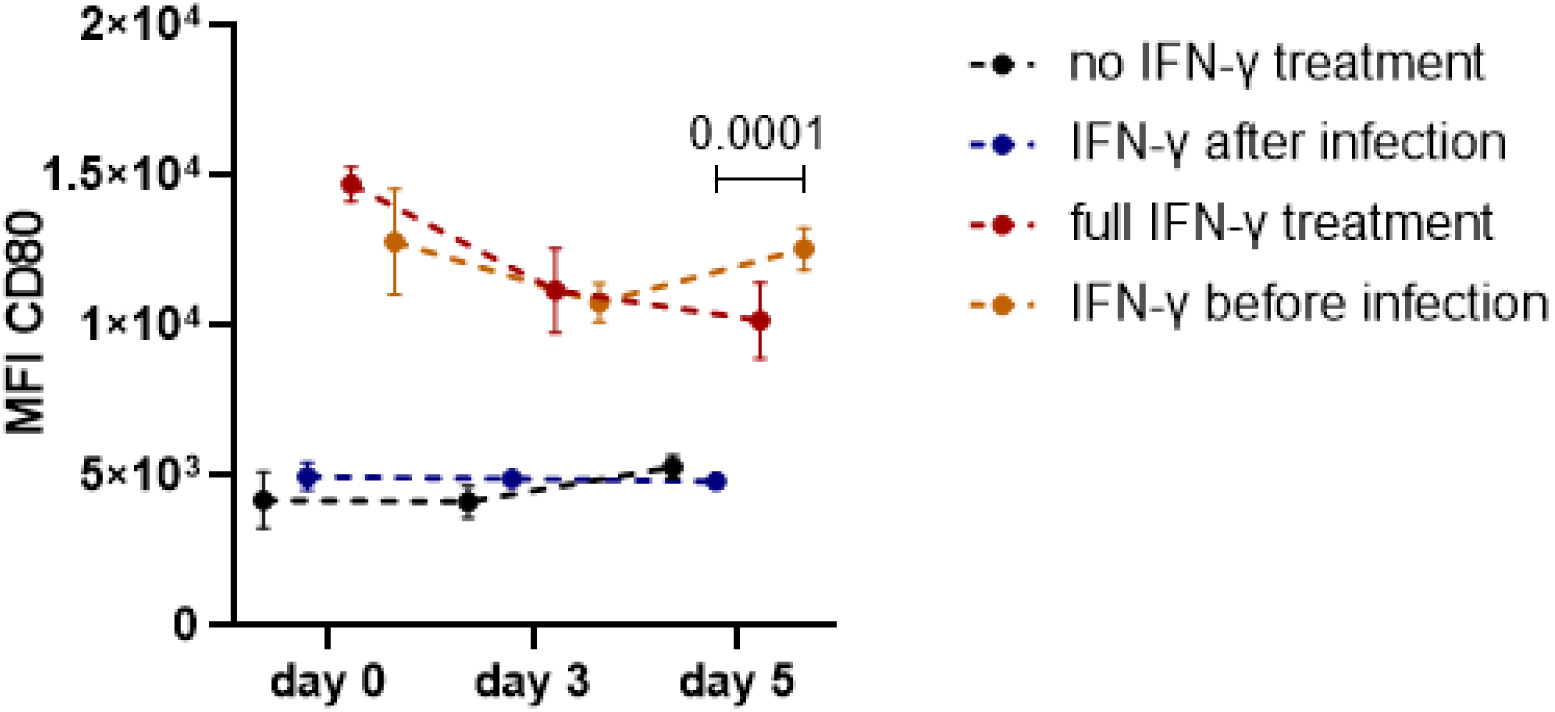
IFN-γ activation before infection enhances CD80 expression in BMDMs. Median fluorescence intensity (MFI) of CD80 in infected BMDMs was measured by flow cytometry at days 0, 3 and 5 post-infection. Data are shown as mean ± SD (n = 3). *P* values were determined by unpaired two-tailed Student’s t-test (before vs. after infection) and corrected for multiple testing using the two-stage linear step-up method of Benjamini–Krieger–Yekutieli.

**Supplementary Figure 5.**
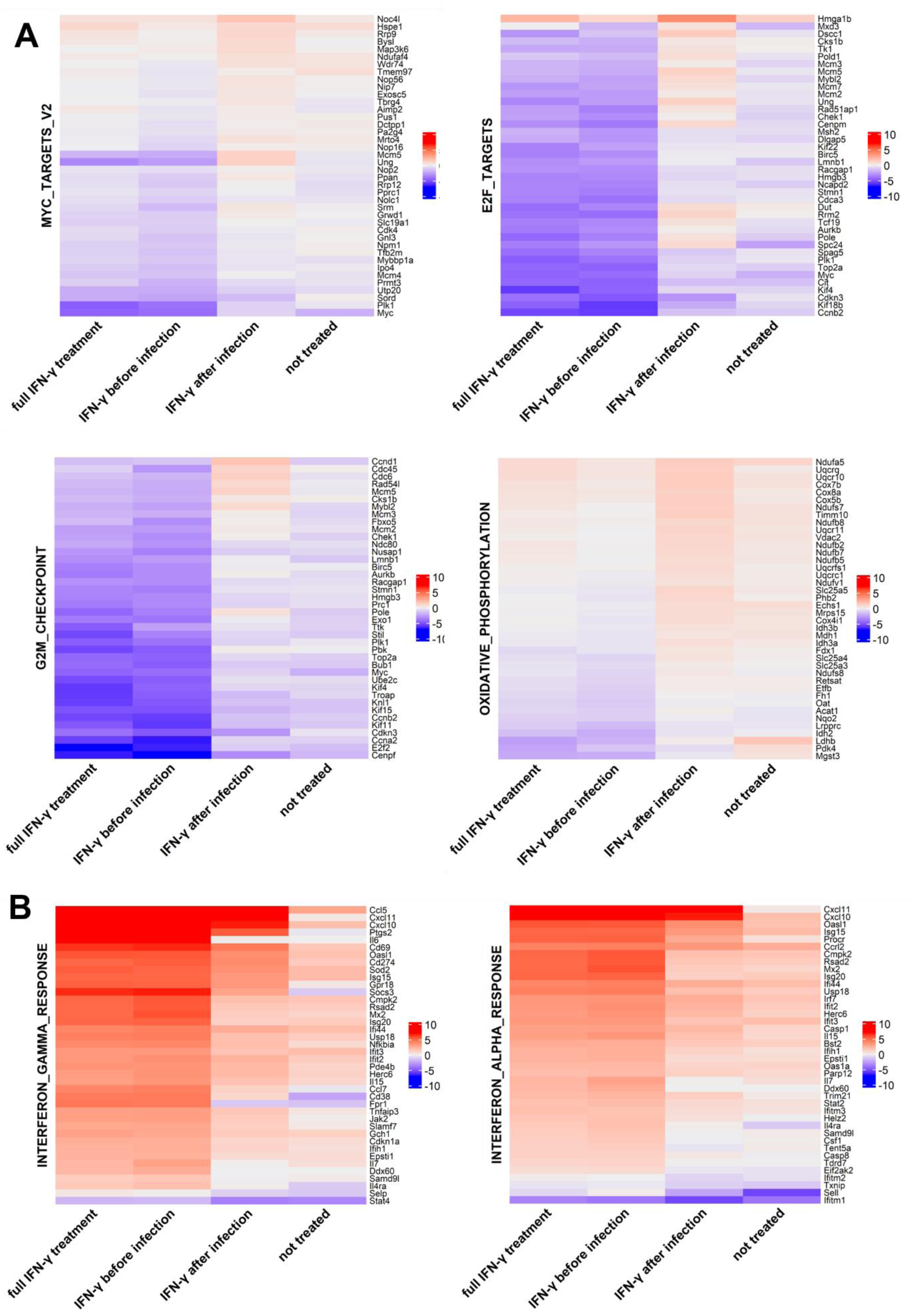
Distinct transcriptional profiles in BMDMs depending on IFN-γ activation. Heatmaps of leading-edge genes (top 40 genes) of downregulated **A)** and upregulated **B)** Hallmark gene sets across all four conditions 24 h after infection. The differential expression is calculated as mean log_2_ fold-change per condition compared to untreated, uninfected BMDMs (n = 3)

**Supplementary Figure 6.**
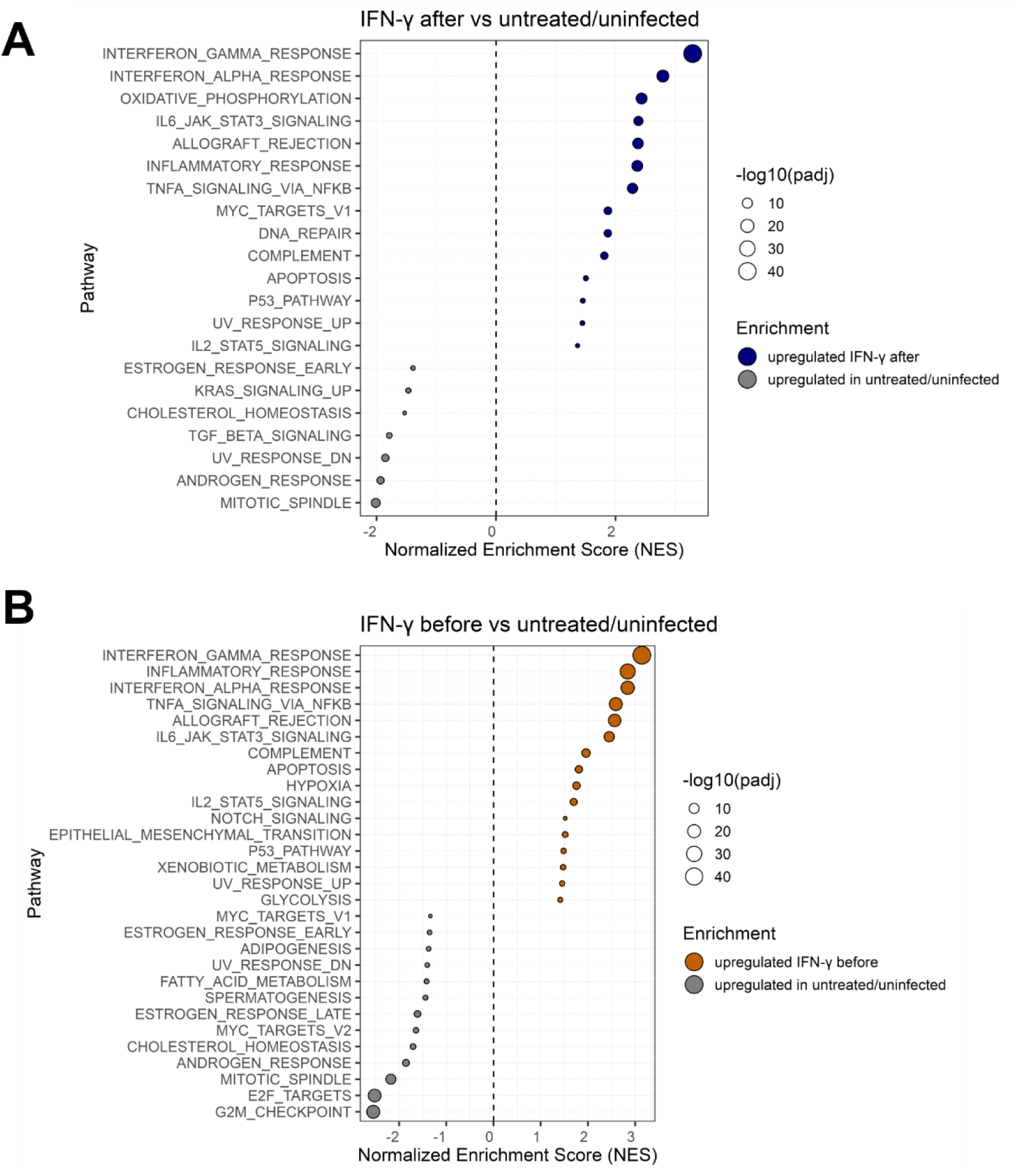
Gene set enrichment analysis of IFN-γ-treated BMDMs. Dot plot of Hallmark pathways enriched in BMDMs treated with **A)** IFN-γ administered after infection versus untreated, uninfected BMDMs and **B)** IFN-γ before after infection versus untreated, uninfected BMDMs 24 h post-infection. Dots indicate normalized enrichment scores (adjusted *P* < 0.05), with positive values indicating enrichment in the IFN-γ-treated condition. NES values were computed by fgsea on MSigDB Hallmarks (v2024.1.Mm).

**Supplementary Figure 7.**
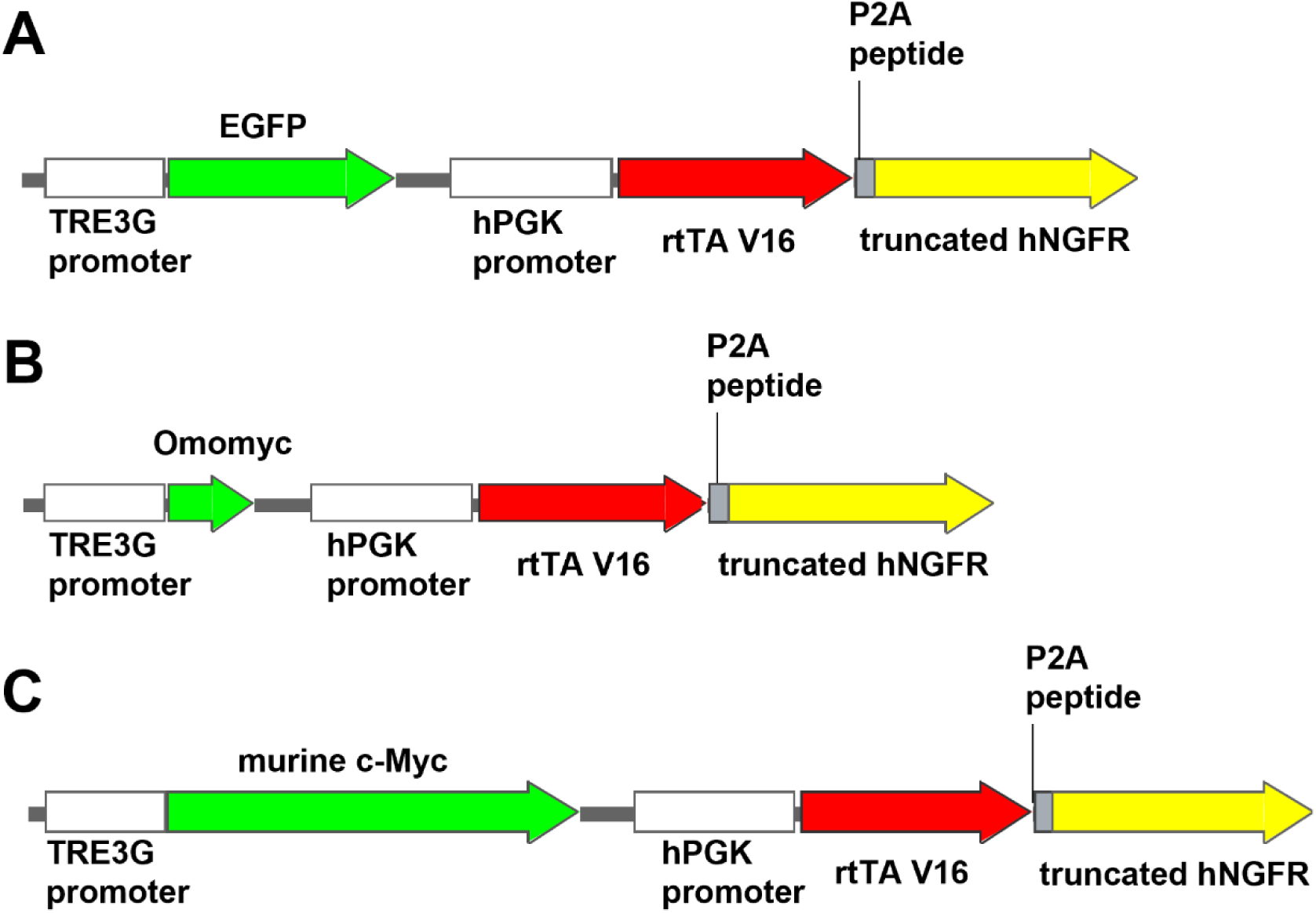
Structure of lentiviral constructs containing the Tet-on system. Structure of the Tet-on constructs expressing different genes: **A)** enhanced green fluorescent protein (EGFP) **B)** Omomyc. **C)** murine c-Myc. The Human phosphoglycerate kinase 1 promoter (hPGK) is constitutively active in most human and mouse cell types. It regulates the transcription of the rtTAV16-P2A-NGFR construct. rtTAV16 is an improved version of reverse tetracycline-controlled transactivator effective at low doses of doxycyline (100ng/mL). Truncated human nerve growth factor receptor (NGFR) is the extracellular domain of the human NGFR, used in this study as marker of transduction. Porcine teschovirus 2A peptide (P2A) is a self-cleaving peptide that allows the production of two separate proteins from one transcript^75^.

**Supplementary Figure 8.**
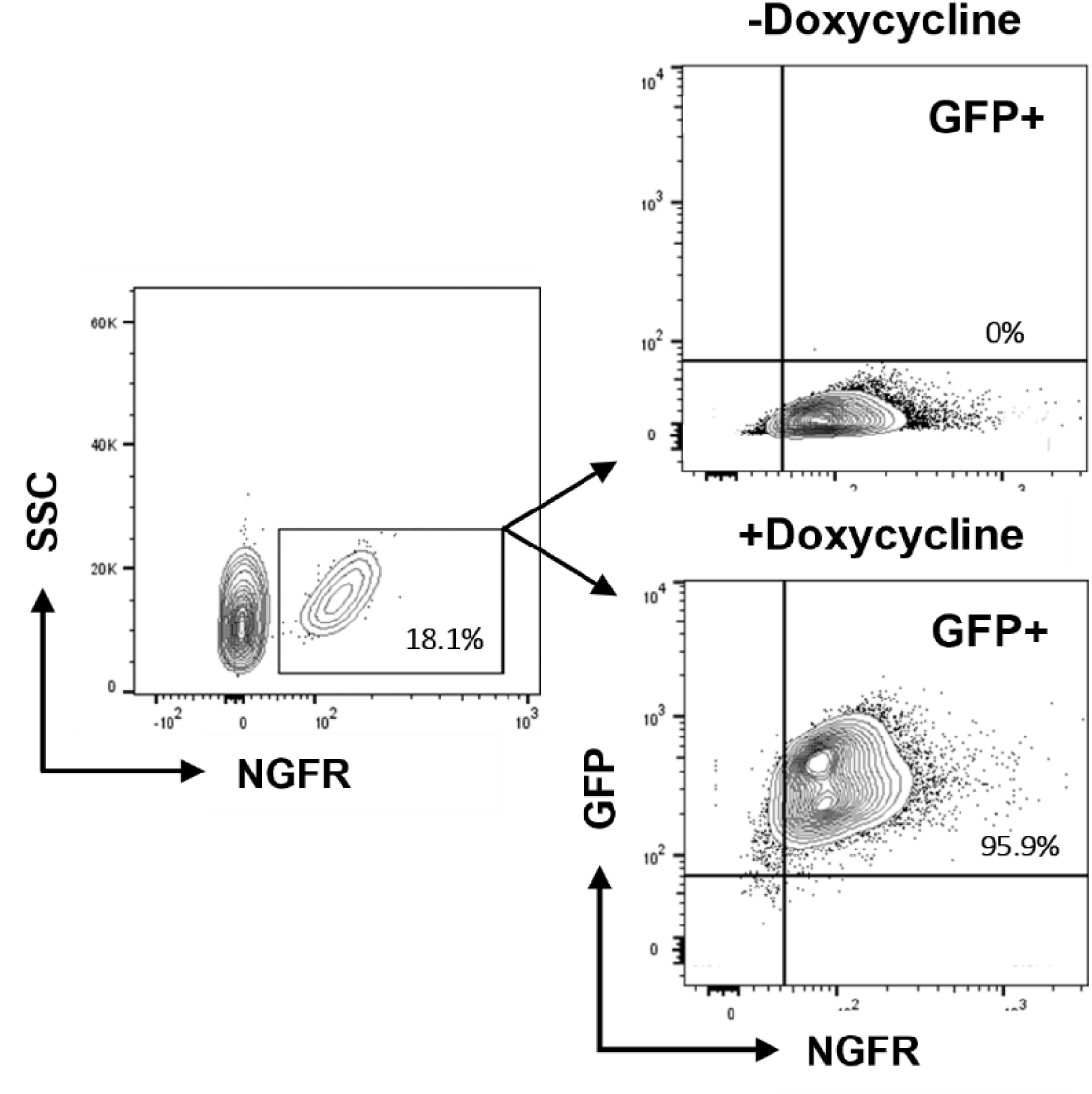
Doxycycline-inducible GFP expression in transduced BMDMs. Representative flow cytometry plots of BMDMs transduced with lentivirus expressing Tet-on regulated GFP, with or without doxycycline (100ng/mL). Bone marrow cells were sorted 24 h after transduction based on the expression of the transduction reporter NGFR. After 6 days, cells were exposed to doxycycline for 24 h and assessed using flow cytometry.

**Supplementary Figure 9.**
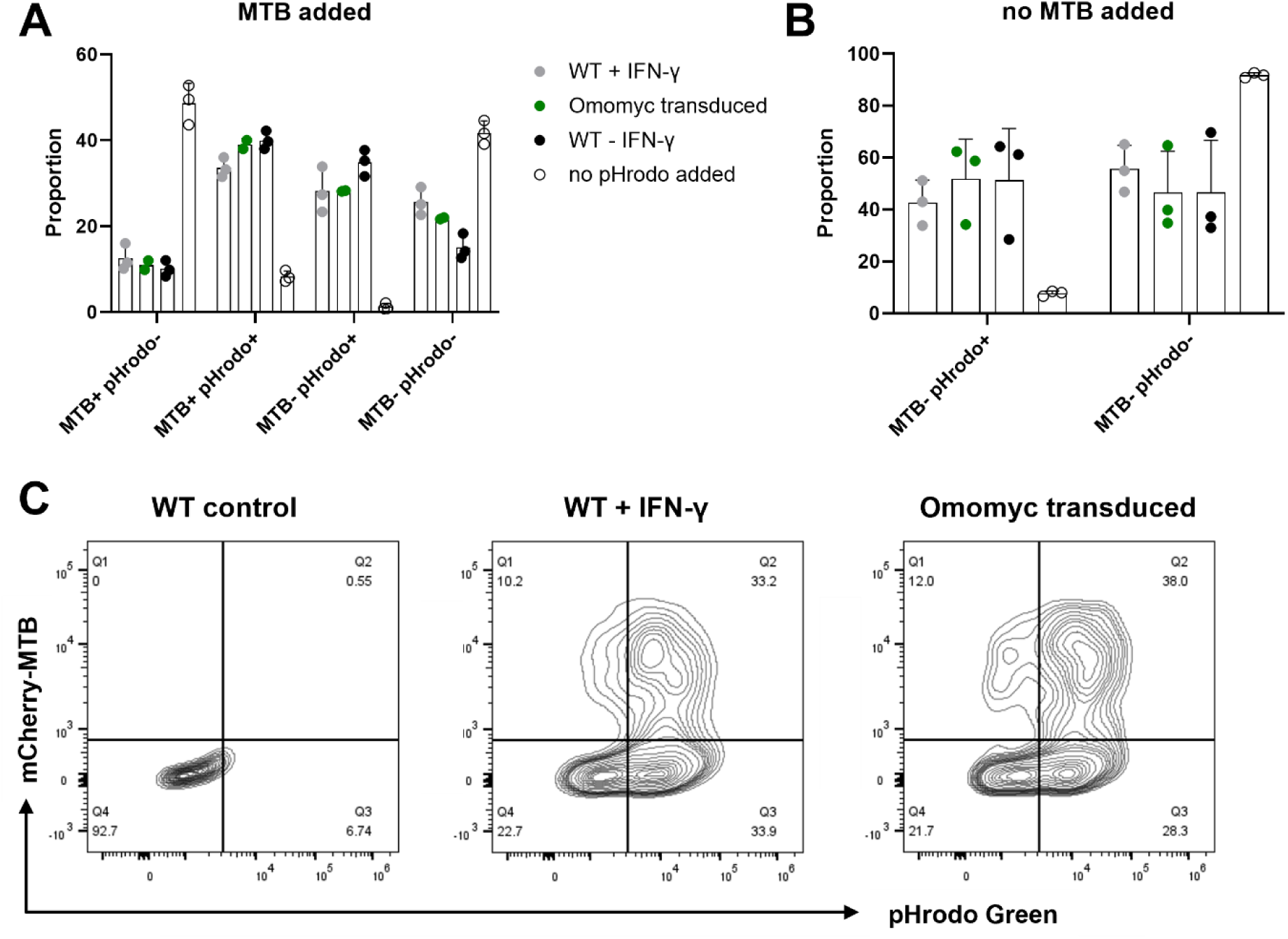
c-Myc perturbation does not alter BMDM phagocytosis or phagosomal acidification. **A)** Quantification of the proportion of BMDMs that have internalized heat-killed, mCherry-expressing MTB and that exhibit an acidic phagosomal compartment (pHrodo-labelled bioparticles) across all experimental conditions, as assessed by flow cytometry (n = 3). **B)** Proportion of BMDMs with acidic phagosomes (pHrodo⁺) in each condition, measured by flow cytometry (n = 3). **C)** Representative flow-cytometry plots showing gating for mCherry⁺ (phagocytosis) and pHrodo⁺ (acidification) populations.

**Supplementary Figure 10.**
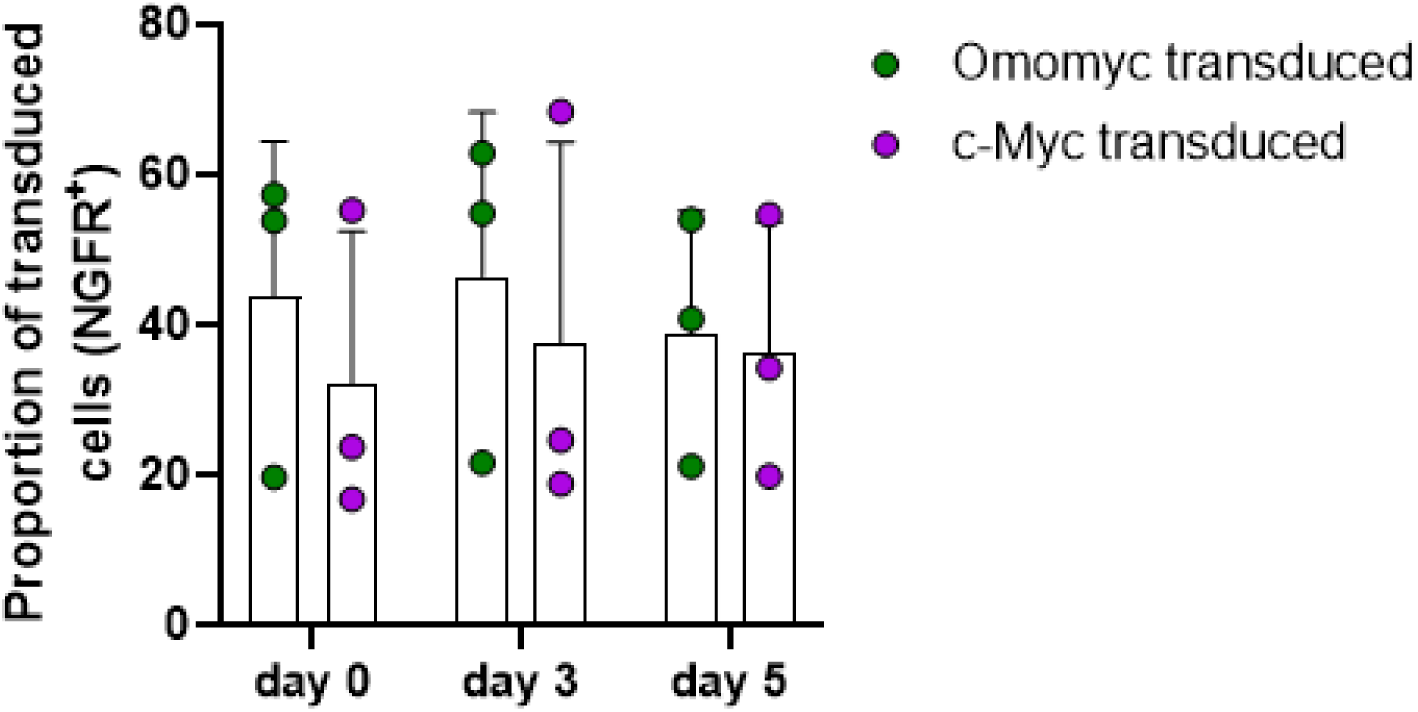
Transduction efficiency of BMDMs across independent experiments. Scatter plot showing the percentage of NGFR^+^ BMDMs, used as reporter for transduction, measured by flow cytometry at each timepoint. BMDMs were lentivirally transduced with either Omomyc or c-Myc constructs. Each data point represents the mean proportion of NGFR^+^ cells from an independent experiment (3 independent experiments).

**Supplementary Figure 11.**
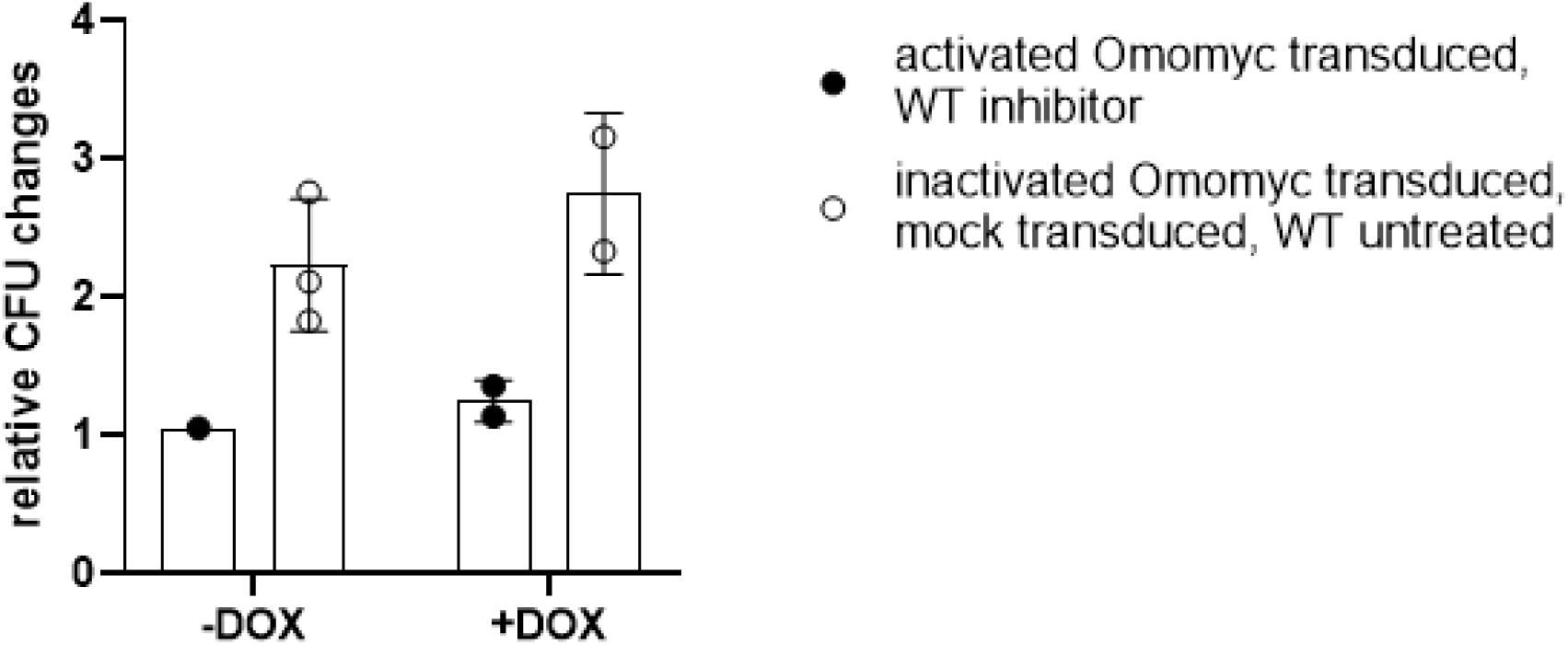
Presence of doxycycline does not affect bacterial growth in CFU assays. Bar plot of relative changes in CFU at day 3 (normalized to day 0) post-infection in the presence or absence of doxycycline (100 ng/mL). Filled and open circles indicate conditions with expected inhibitor-induced kinetics versus those exhibiting standard CFU kinetics, respectively.

**Supplementary Figure 12.**
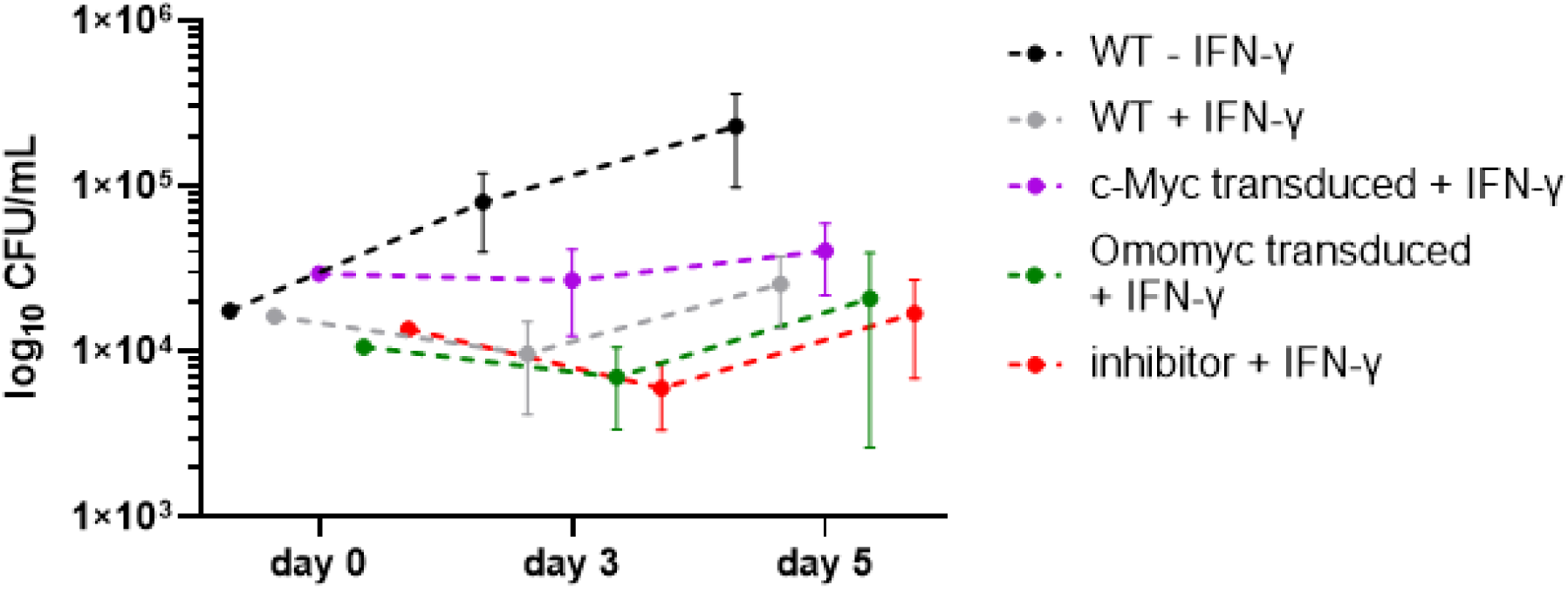
Transduced and c-Myc perturbed BMDMs show comparable CFU growth to IFN-γ activation. Quantification in colony forming units per milliliter (CFU/mL) of intracellular MTB at day 0 (immediately after phagocytosis) and days 3 and 5 post-infection in IFN-γ–activated WT, c-Myc transduced, Omomyc transduced, and inhibitor-exposed BMDMs.

**Supplementary Figure 13.**
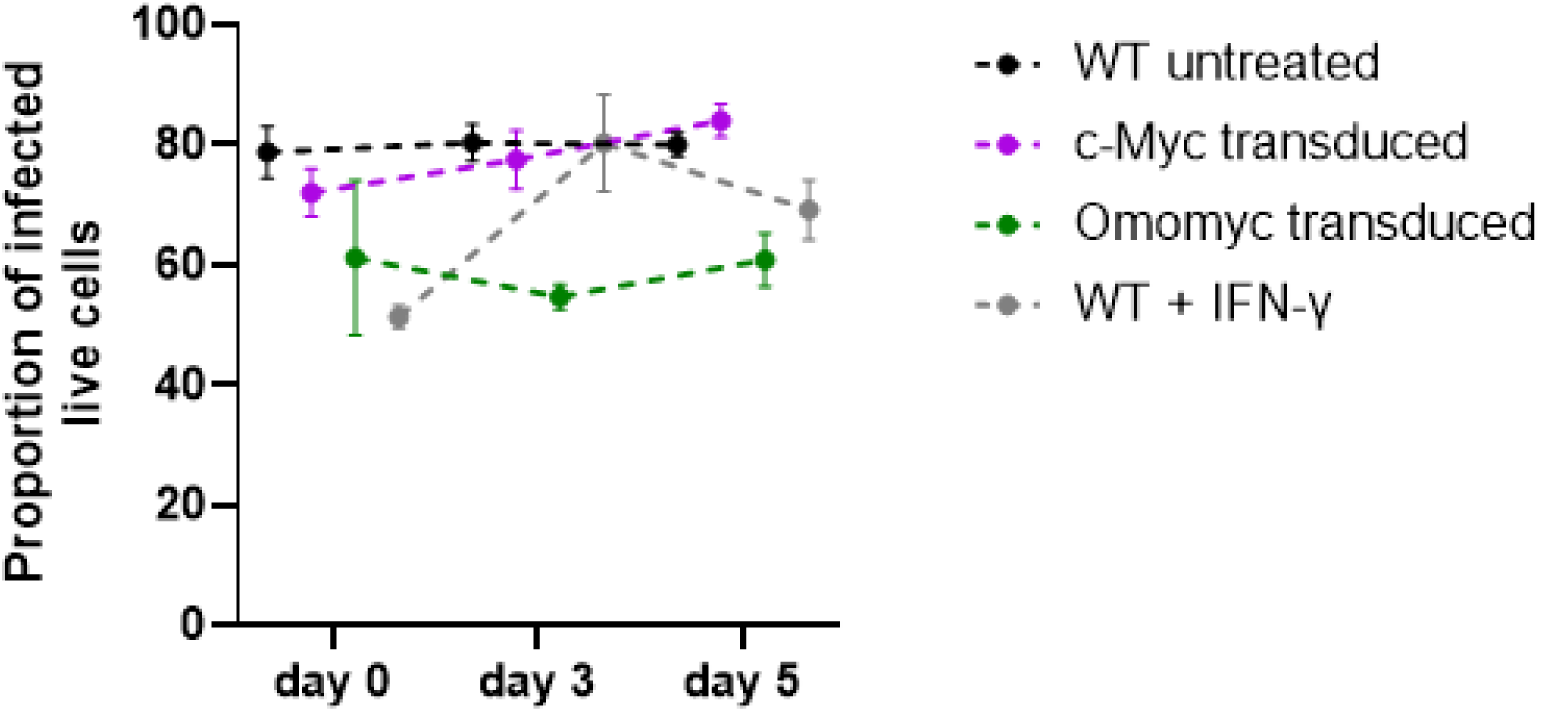
BMDM viability remains stable by Omomyc-mediated c-Myc inhibition over time. Proportion of infected (mCherry^+^) live cells among infected BMDMs in WT, c-Myc–expressing, Omomyc– expressing, or IFN-γ–treated WT conditions, as assessed by flow cytometry. Live cell percentages were measured immediately after phagocytosis (day 0) and at days 3 and 5 post-infection.

**Supplementary Figure 14.**
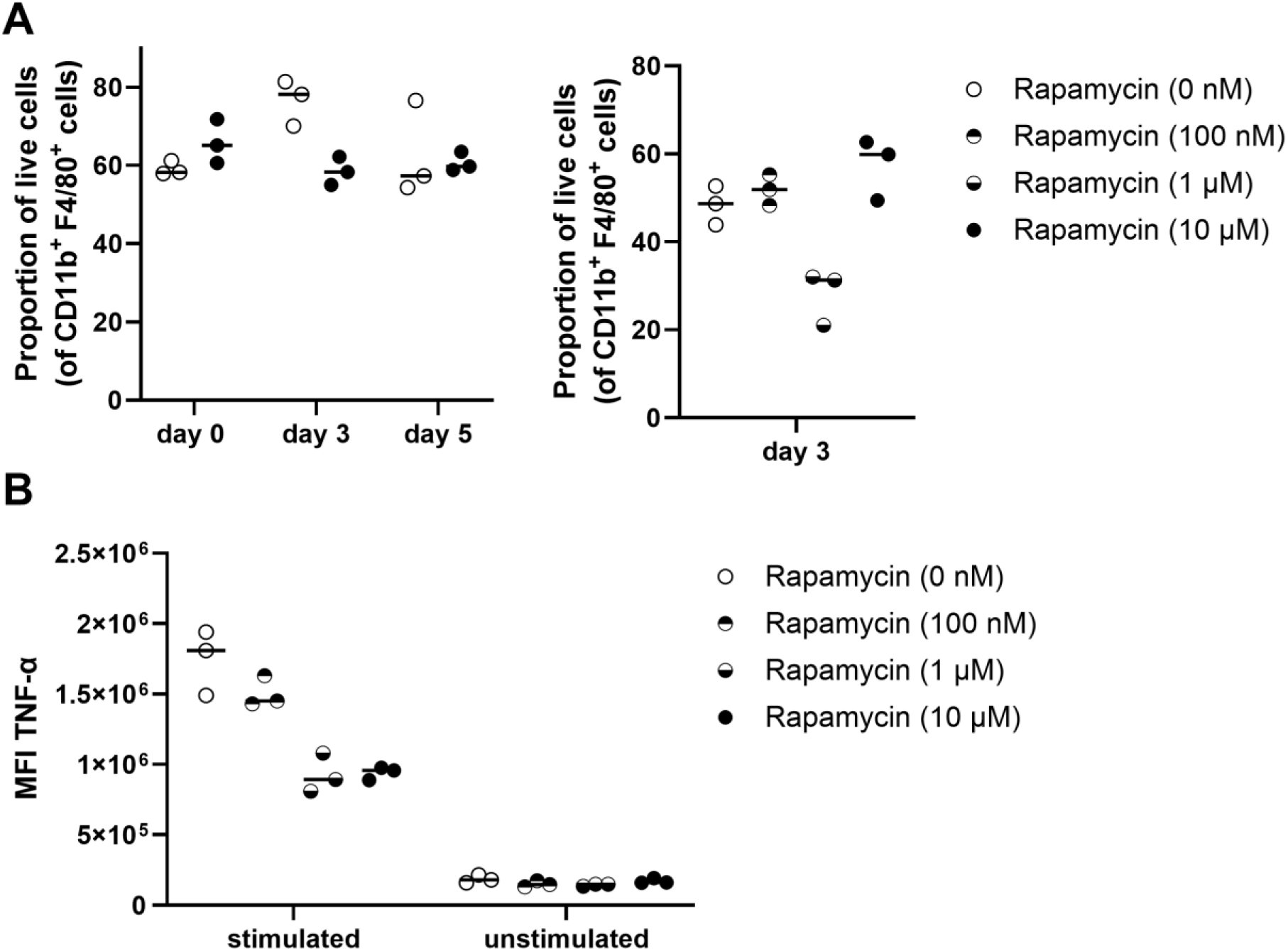
Rapamycin does not substantially influence survival in macrophages. A) Proportion of live CD11b^+^ F4/80^+^ conditionally immortalized macrophages at days 0, 3 and 5 following treatment with increasing concentrations of rapamycin, as assessed by flow cytometry. B) Median fluorescence intensity (MFI) of TNF-α in LPS (10 ng/mL) stimulated for 6 h with Brefeldin A, and unstimulated macrophages, treated with increasing concentrations of rapamycin, measured by flow cytometry.

**Supplementary Figure 15.**
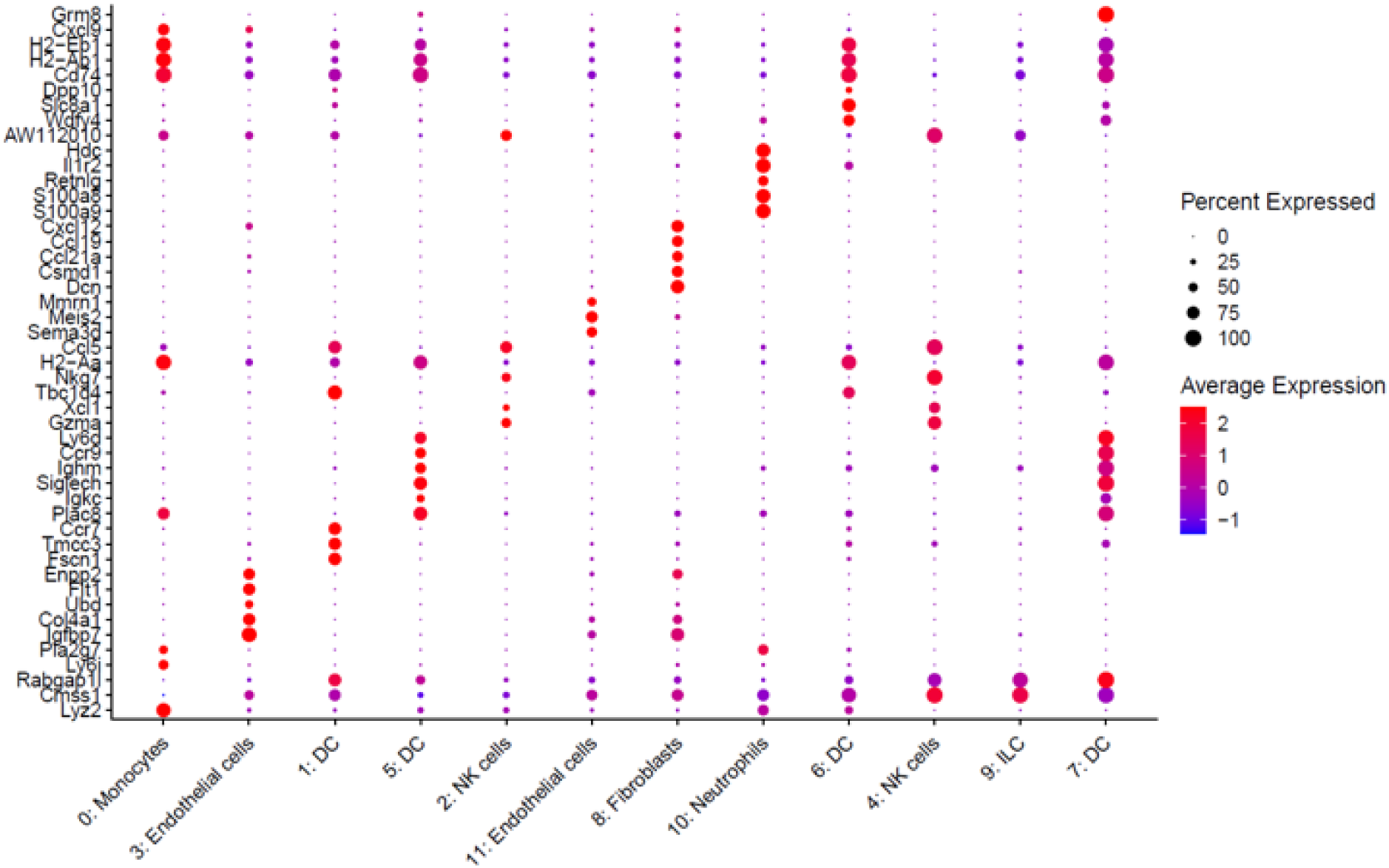
Cluster-defining genes in scRNA-seq. Cluster-defining genes were used to deconvolute scRNA-seq data and generate clusters using the Seurat package^28^.

**Supplementary Figure 16.**
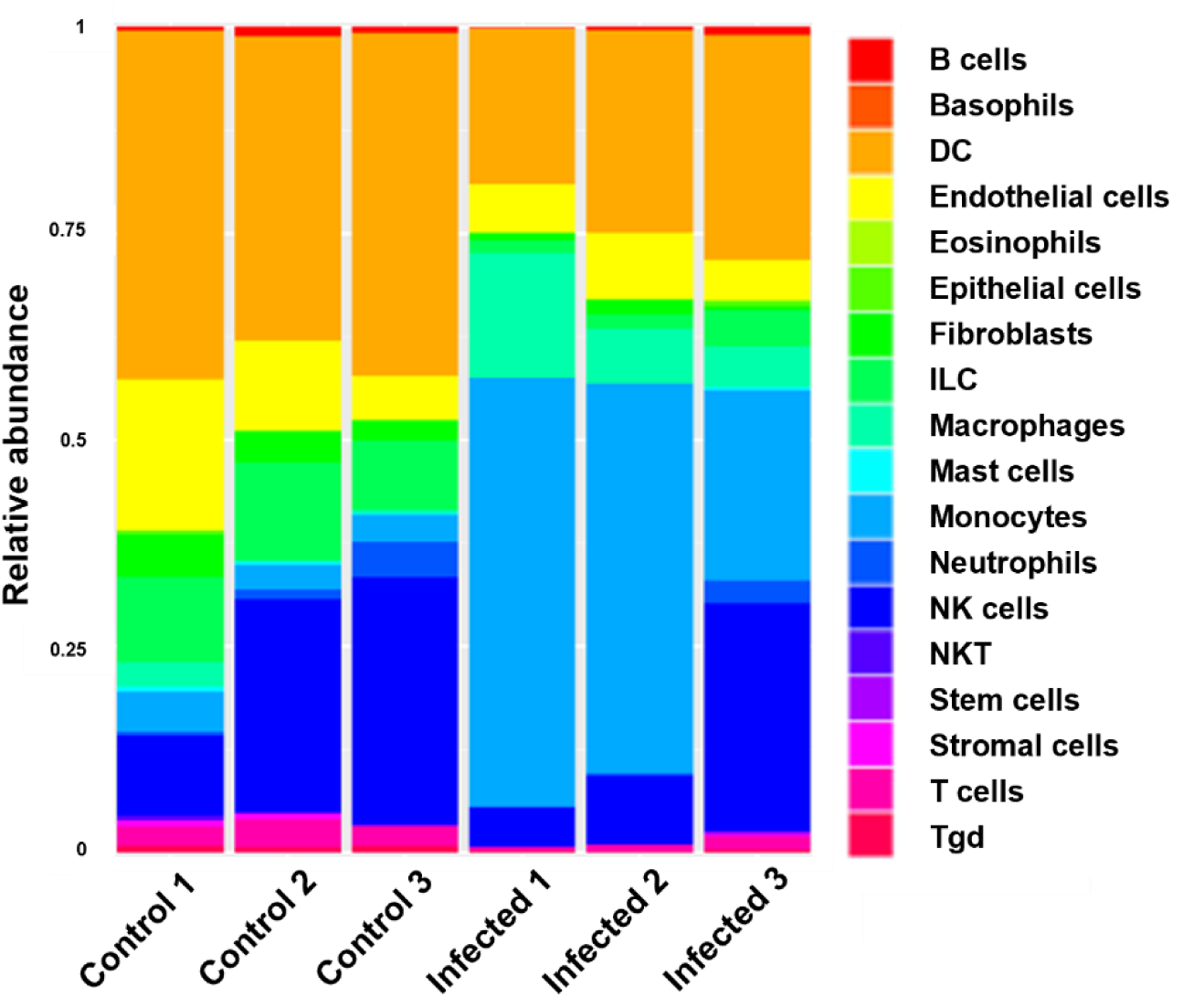
Identification of cell populations by scRNA-seq. Identification of cell populations by scRNA-seq in lymph nodes from mice with contained MTB infection (CMTB) and uninfected controls. Relative abundance of each cell population is shown; note that T and B cells were depleted before sequencing and therefore appear at very low frequency.

**Supplementary Table 1.**
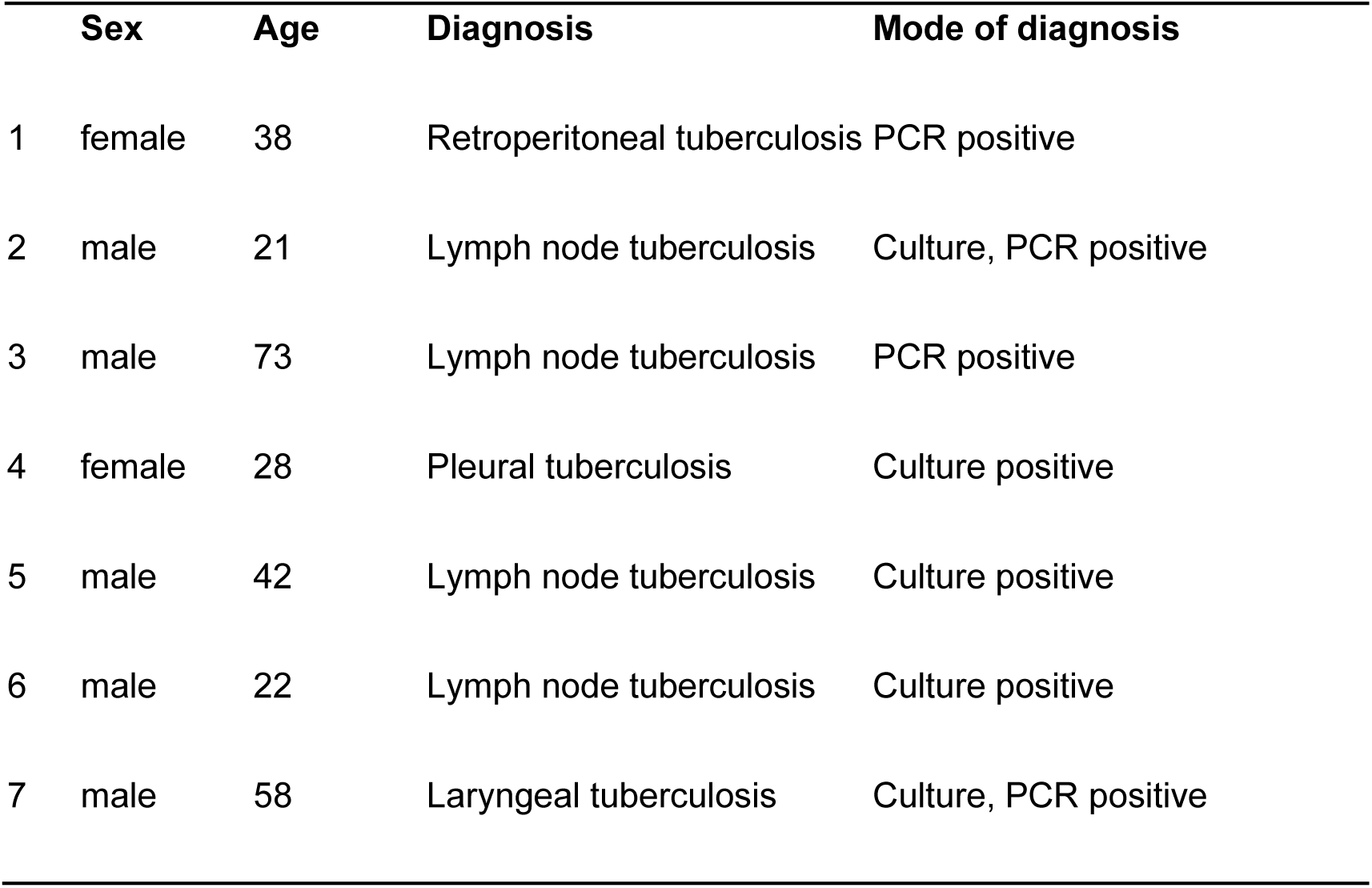
Patient characteristics.

